# Descending control and regulation of spontaneous flight turns in *Drosophila*

**DOI:** 10.1101/2023.09.06.555791

**Authors:** Ivo G. Ros, Jaison J. Omoto, Michael H. Dickinson

**Affiliations:** Division of Biology and Bioengineering California Institute of Technology 1200 E. California Blvd. Pasadena CA, 91125, USA

**Keywords:** Flight saccades, command neurons, spontaneous activity, local search, long-distance dispersal

## Abstract

The clumped distribution of resources in the world has influenced the pattern of foraging behavior since the origins of life, selecting for a common locomotor search motif in which straight movements through resource-poor regions alternate with zig-zag exploration in resource-rich domains (Berg, 2000). For example, flies execute rapid changes in flight heading called body saccades during local search (Censi et al., 2013; Collett and Land, 1975; Schilstra and van Hateren, 1999; Wagner and Land, 1986), but suppress these turns during long-distance dispersal (Giraldo et al., 2018; Leitch et al., 2021) or when surging upwind after encountering an attractive odor plume (Budick and Dickinson, 2006; van Breugel and Dickinson, 2014). Here, we describe the key cellular components of a neural network in flies that generates spontaneous turns as well as a specialized neuron that inhibits the network to promote straight flight. Using 2-photon imaging, optogenetic activation, and genetic ablation, we show that only four descending neurons appear sufficient to generate the descending commands to execute flight saccades. The network is organized into two functional couplets—one for right turns and one for left—with each couplet consisting of an excitatory (DNae014) and inhibitory (DNb01) neuron that project to the flight motor neuropil within the ventral nerve cord. Using resources from recently published connectomes of the fly brain (Scheffer et al., 2020; Dorkenwald et al., 2023; Schlegel et al., 2023), we identified a large, unique interneuron (VES041) that forms inhibitory connections to all four saccade command neurons and created specific genetic driver lines for this cell. As predicted by its connectivity, activation of VES041 strongly suppresses saccades, suggesting that it regulates the transition between local search and long-distance dispersal. These results thus identify the critical elements of a network that not only structures the locomotor behavior of flies, but may also play a crucial role in their foraging ecology.

## Introduction

When searching their environment, many species of flies execute a series of straight flight segments interspersed with extremely rapid turns lasting less than ∼50 ms. These brief turns were termed saccades, in analogy with the rapid eye movements of primates (Collett and Land, 1975). In flies, however, the entire body quickly rotates during each saccade, not just the eyes. These saccades can benefit the animals in several ways. From a sensory perspective, saccades may restrict the deleterious effects of motion blur to brief moments interjected within longer sequences of gaze stabilization (Cellini et al., 2021; Land, 1999; Schilstra and van Hateren, 1998). Brief bursts of saccades in the same direction may aid local search strategy by allowing the animal to quickly scan the local environment for salient visual and olfactory features (Heisenberg and Wolf, 1984). More recently, it has been suggested that comparing sensory measurements before and after each saccade might enable flies to estimate key parameters that are otherwise not directly measurable, such as the direction and magnitude of the ambient wind (van Breugel et al., 2022). For all these hypotheses, the timing between saccades is critical, and several factors are thought to influence the frequency of the spontaneous turns. For example, tethered flies flying in a closed-loop configuration will greatly reduce the saccade rate and instead actively steer toward salient visual objects such as vertical edges—a robust fixation behavior that they maintain for several hours (Götz, 1987). Similarly, flies also reduce saccade rates in the presence of a small bright spot, which they presumably interpret as the sun. In this case, however, they do not necessarily steer directly toward the distant bright object, but rather maintain it at an arbitrary azimuthal angle (Giraldo et al., 2018). Patterns of polarized light that mimic those in the natural sky induce a similar reflex (Warren et al., 2018; Weir and Dickinson, 2012). This orientation behavior using celestial cues, termed menotaxis, may allow flies and other insects (El Jundi et al., 2016) to maintain a constant heading over long distances (<10km) while dispersing (Coyne et al., 1982; Leitch et al., 2021). Flies also suppress saccades in the presence of a constant odor stream to promote their progress upwind toward a food source (Barrows, 1907; Budick and Dickinson, 2006; van Breugel and Dickinson, 2014). Despite its importance with respect to multiscale search, the neural mechanisms that underlie both the generation of saccades and their suppression during dispersal and plume tracking remain largely unknown.

Within the central nervous system of insects, descending neurons (DNs) constitute a critical stage in the transformation of sensory input in the brain into motor commands in the ventral nerve cord (VNC) (Heinrich, 2002). *Drosophila* possess ∼650 pairs of descending neurons (Cheong et al., 2023; Hsu and Bhandawat, 2016; Namiki et al., 2018), some of which appear to function as specialized command neurons for specific behaviors including courtship (Kohatsu et al., 2011; von Philipsborn et al., 2011), walking backward (Bidaye et al., 2014), turning (Rayshubskiy et al., 2020), take off (von Reyn et al., 2014; Wyman et al., 1984), and landing (Ache et al., 2019). Thus, DNs provide a logical starting point for investigating the circuits that generate and regulate flight saccades. A prior study using electrophysiology and 2-photon imaging implicated a pair of DNs, called AX, in the execution of spontaneous and visually-elicited flight saccades (Schnell et al., 2017). However, this work employed a broad GAL4-driver line (R65G08) and a separate study found no significant effect on saccade rate when the cells within this line were silenced via a temperature-sensitive dynamin mutation that blocks synaptic transmission (Ferris et al., 2018). To confirm and further investigate the role of AX in saccade generation, we constructed a sparse split-GAL4 driver for this neuron better suited for functional imaging, optogenetic activation, and genetic ablation. From here on, we will refer to the AX neuron as DNae014, as this is its identifier in the FlyWire database (Dorkenwald et al., 2023; Schlegel et al., 2023). Because we suspected that multiple pairs of DNs might be involved in driving the bilateral wing movements exhibited during saccades, we screened a large collection of split-GAL4 lines (Namiki et al., 2018) for additional cells that function to trigger rapid turns. Our screen led to the discovery of one additional cell (DNb01) that together with DNae014 appears to form two functional couplets that drive saccades to the left and right. We systematically analyzed the upstream inputs to these DNs using emerging connectomes of the *Drosophila* brain (Scheffer et al., 2020; Dorkenwald et al., 2022; Dorkenwald et al., 2023; Schlegel et al., 2023) and found a large, unique inhibitory interneuron (VES041) whose activation suppresses saccades thereby promoting straight flight. Thus, we have identified the core cellular components of a network that generates and regulates spontaneous flight maneuvers in flying *Drosophila*.

## Results

To confirm the role of DNae014 in generating flight saccades, we created a split-GAL4 line (Luan et al., 2020) that selectively targets this neuron for functional imaging, optogenetic activation and silencing, and genetic ablation. To characterize the activity of individual DNae014 neurons, we used 2-photon functional imaging to record from DNae014 cells on the left and right side of the brain, while simultaneously tracking wing motion during flight (Fig. 1). The difference in GCaMP7f fluorescence between the bilateral pair of cells (R-L ΔF/F) correlated remarkable closely with the difference in wingstroke amplitude (L-R WSA) (Fig. 1f, h). We used a simple classifier to determine the timing of saccades based on transient differences in WSA (dots in Fig. 1h, i). We interpreted these saccades as spontaneous events because our experiments were conducted in darkness. In addition to DNae014, we identified another cell, DNb01, whose activity is also tightly correlated with the production of spontaneous saccades (Fig.1d, e, g, i). Across individuals, the bilateral difference in cell fluorescence (R-L ΔF/F) explained 89±4% (for DNae014) and 90±3% (for DNb01) of the variance in differential wingstroke amplitude (L-R) (Fig. 1k). The activity levels between the left and right DNae014 cells followed an inverse, highly non-linear relationship, analogous to ‘flip-flop’ components in digital electronic circuits (Fig. 1j), and reminiscent of neurons identified in the steering behavior of male silkmoths (Olberg, 1983). The relationship for the two DNb01 cells was also inverse, but much more linear. Given the complex relationship between the electrical activity of a neuron and its GCaMP signal, these subtle differences in the bilateral patterns of the DNae014 and DNb01 cells may not be functionally relevant; but nevertheless, the activities of both neurons correlate remarkably well with ipsilaterally directed saccades (Fig. 1l).

**Fig. 1.**
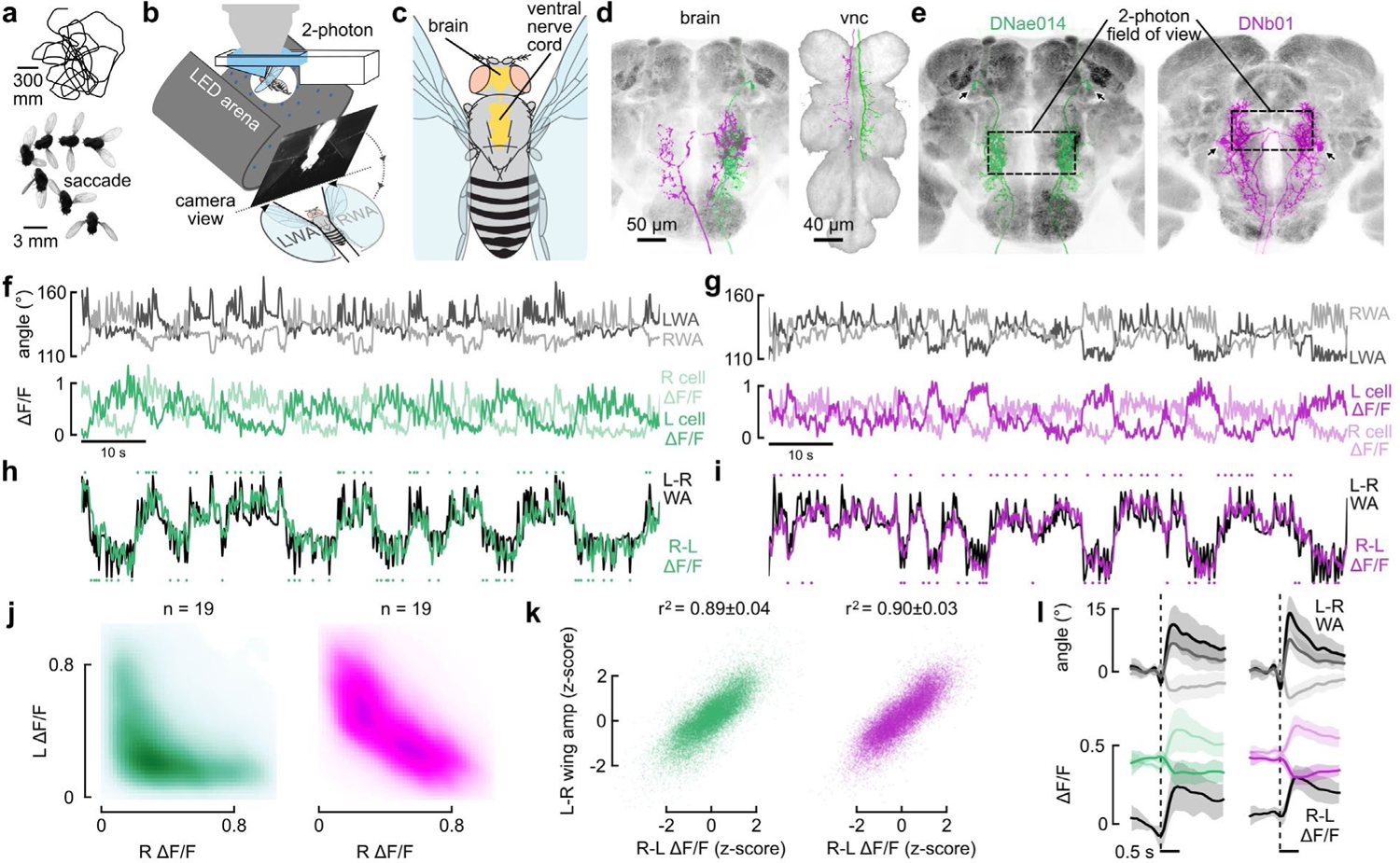
Activities of DNae014 and DNb01 correlate strongly with spontaneous saccades. **a**, Flight trajectory and photomontage illustrating a single saccade, adapted from (Censi et al., 2013; Muijres et al., 2014). **b**, Schematic of set-up for monitoring neural activity with GCaMP7f and simultaneously tracking wingstroke amplitudes during flight. **c,** Fly with central nervous system. **d**, Anatomy of right DNae014 (green) and DNb01 (magenta) cells in the brain and VNC. **e**, Bilateral expression patterns of DNae014 and DNb01 cells in the brain from split-GAL4 lines, showing the fields of view scanned in 2-photon experiments (dashed boxes). Black arrows indicate somata. **f**, Top: left (dark gray) and right (light gray) wingstroke amplitudes. Bottom: normalized GCaMP7f fluorescence (ΔF/F) in left and right DNae014 cells during a 90 s flight epoch. **g**, Similar to **f**, but for DNb01. **h**, Right-left difference in DNae014 signals (green) superimposed with left-right difference in wingstroke amplitudes (black). Both traces are normalized by z-score from the traces above in **f**. Automatically detected right and left saccades depicted as green dots above and below, respectively. **i**, Similar to **h**, but for DNb01. **j**, Pooled regressions for DNae014 (green) and DNb01 (magenta) of left vs. right cell activity (ΔF/F), normalized by z-score (n = 19 flies) and plotted as a kernel density estimate. **k**, Regressions of L-R wingstroke amplitude vs. R-L ΔF/F, across 19 flies for DNae014 (green; r^2^ = 0.89±0.04, mean±sd) and DNb01 (magenta, r^2^ = 0.90±0.03, mean±sd). **l**, Changes in DNae014 (left column) and DNb01 (right column) cell activity aligned to the onset of spontaneous flight saccades. Top traces: ΔF/F signals for right cell (light color), left cell (dark color), and right-left difference (black). Bottom traces: baseline-subtracted wingstroke amplitudes of left (dark gray), right (light gray) wings, along with left-right difference (black). Solid lines indicate mean of means for all individuals (n = 19 flies for each dataset); shaded areas indicate boot-strapped 95% CIs.

Despite similar activity patterns, DNae014 and DNb01 are morphologically distinct. DNae014 possesses smooth, dendritic processes in the lateral accessory lobe (LAL) and superior posterior slope (SPS) with varicose terminals in the gnathal ganglion (GNG) and wing neuropil of the VNC (Fig. 1c, d). As described before based on dye fills following whole cell recordings (Schnell et al., 2017), all processes of the DNae014 cell are ipsilateral to the soma in both the brain and VNC. In contrast, DNb01 crosses the midline via the LAL commissure with varicose terminals in the contralateral SPS, posterior ventrolateral protocerebrum (PVLP) and LAL, and sends a descending axon to the wing neuropil of the VNC contralateral to the cell body (Fig. 1d, e). Both cells arborize extensively within the LAL, a region implicated in turning behaviors in a variety of insect species (Homberg, 1994; Steinbeck et al., 2020). Although we did not record from DNae014 and DNb01 simultaneously, transient changes in the activity of either corresponds with a rapid alteration in wing motion, and conversely, every change in wing motion corresponds with transients in the activity of the neurons. Based on this close correlation (Fig. 1k), we tentatively conclude that these two pairs of DNs are co-active during flight saccades (Fig. 1).

To confirm the prior observation that DNae014 responds to expanding visual patterns and test whether DNb01 exhibits a similar phenomenology, we imaged bilateral activity of the DNs while presenting a looming stimulus at different azimuthal positions (Extended Data Fig. 2). For both DNae014 and DNb01, a looming spot presented in one visual hemisphere increased activity in cells on the opposite side (Extended Data Fig. 2b). Although the absolute magnitude of the measured ΔF/F response for DNae014 was larger, this result based on Ca^2+^ imaging need not reflect an actual difference in the membrane-level responses of the two DNs. Recently, other descending neurons have been identified that appear to mediate looming responses during flight (Kim et al., 2023), which suggests that DNs other than DNae014 and DNb01 may be involved in visually elicited avoidance reactions.

To test whether DNae014 and DNb01 are sufficient to elicit saccades, we used several methods to activate individual neurons on either the left or right side of the brain while measuring the resulting changes in wing motion. First, we drove CsChrimson bilaterally in DNae014 cells, but used focal 2-photon excitation to unilaterally activate neurons on one side of the brain or the other (Fig. 2a-c). Activation of the left DNae014 cell reliably elicited a saccade to the left and activation of the right cell elicited a saccade to the right, consistent with the correlation recorded during spontaneous activity (Fig. 1). These evoked syndirectional saccades also confirm prior experiments that used focal injection of ATP to activate ectopically expressed P_2_X_2_ channels (Lima and Miesenböck, 2005) in these neurons (Schnell et al., 2017). Focal 2-photon CsChrimson activation was not feasible for single DNb01 cells, because each neuron branches extensively in both the left and right hemispheres (Fig. 1d, e). Instead, we unilaterally activated the DNs using the SPARC system (Isaacman-Beck et al., 2020) to express CsChrimson (Fig. 2d). Because the SPARC reagent produced flies with CsChrimson in either both targeted cells, neither cells, the left cell only, or the right cell only, we scored the expression pattern of each individual after testing their behavioral responses to pulses of 617 nm light. A distinct advantage of the SPARC method is that it enforces a blinded experimental design, with the subset of flies in which neither of the targeted cells expressed CsChrimson additionally serving as a convenient control group. Unilateral activation of DNae014 evoked the expected changes in wing motion based on our 2-photon experiments (Fig. 2g), and unilateral activation of DNb01 also evoked syndirectional saccades. However, the saccades evoked by DNb01 activation were smaller in amplitude and longer in time course than those evoked by DNae014 activation, suggesting that the two cells drive different components of the saccade motor program within the VNC.

**Fig. 2.**
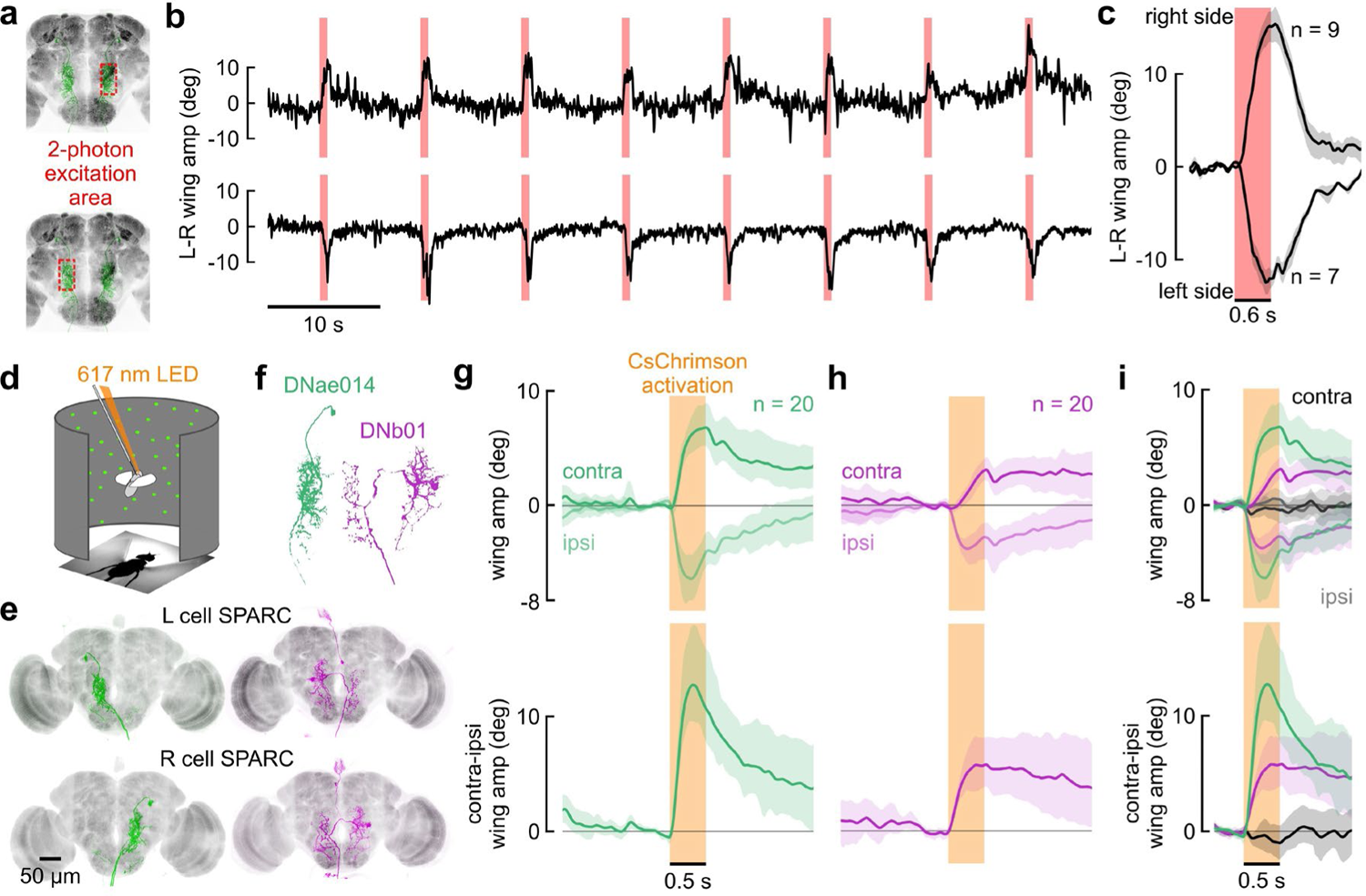
Unilateral activation of either DNae014 and DNb01 elicits directional saccades. **a-c**, Unilateral 2-photon excitation of either left or right DNae014 neurons. **a**, Approximate 2-photon excitation areas of the right and left dendritic arbors of DNae014. **b**, Example traces of L-R wingstroke amplitude during unilateral CsChrimson excitation on the left (top) or right (bottom) side of brain. **c**, L-R wing amplitudes aligned to the excitation light pulse. As in all panels, solid lines indicate the mean of all individual means, the shaded patch indicates the boot-strapped 95% CI; n = 9 flies for right stimulations, n = 7 flies for left stimulations. **d-i**, Unilateral activation of DNae014 or DNb01 neurons in rigidly tethered flies using split-GAL4 and SPARC expression of CsChrimson. **d**, Flight arena with 617 nm excitation light. **e**, Maximum intensity projection of TdTomato signal in examples of unilateral cell expression using SPARC with split-GAL4 drivers for DNae014 (green) and DNb01 (magenta). Nc82 staining shown in grey. **f,** Representative morphology of right DNae014 and DNb01 cells. **g,** Contralateral and ipsilateral wingstroke amplitudes (top) and contralateral-ipsilateral difference (bottom), aligned to activation pulse; n= 20 flies. **h**, Similar to **g**, but for DNb01; n = 20 flies. **i**, Combined data from **g** and **h**, including control flies (black traces, gray envelopes) in which neither cell expressed CsChrimson; n = 20, 20, and 19 for DNae014, DNb01, and controls, respectively.

To assess the necessity of the two pairs of DNs to generate spontaneous saccades, we silenced the neurons via expression of the light-gated GtACR1 channel. We periodically exposed rigidly tethered flies expressing GtACR1 (Govorunova et al., 2015) in DNae014 to 530 nm light pulses (Extended Data Fig. 3a, b). Although the experimental group exhibited a strong decrease in saccade rate when exposed to 530 nm light, empty-vector control flies, which did not express GtACR1, exhibited a substantial increase in saccade rate in response to the pulses of green light (Extended Data Fig. 3c, d). Whereas these silencing results support the hypothesis that suppressing DNae014 activity reduces the frequency of spontaneous saccades, the interpretation is complicated by the large increase in saccade rate exhibited by control flies in response to the activation light. For this reason, we adopted an alternative strategy to test the necessity of the two DNs to saccade by genetically ablating the cells through ectopic expression of the pre-apoptotic genes *reaper* and *hid* (Goyal et al., 2000).

We quantified flight saccades in rigidly tethered flies in which either DNae014 or DNb01 was ablated. Empty vector control flies exhibited a saccade frequency of 0.48 ± 0.18 Hz (mean ± S.D.), consistent with prior studies under both free and tethered flight conditions (Bender and Dickinson, 2006; Censi et al., 2013; Schnell et al., 2017). Genetic ablation of DNae014 reduced saccade frequency to 0.09 ± 0.09 Hz (p<0.001, Kruskal-Wallis test followed by post-hoc comparisons with Tukey’s HSD method), whereas ablation of DNb01 reduced the rate to 0.12 ± 0.14 Hz (p<0.001; Fig. 3b, c). Thus, ablation of either DNae014 or DNb01 significantly reduced the rate of spontaneous saccades but did not eliminate them. This result suggests that bilateral pairs of either DNae014 or DNb01 acting alone can still generate partial saccades, albeit at a reduced spontaneous frequency, or alternatively, that spontaneous saccades that persist following ablation of either cell type might be generated by an alternate pathway.

**Fig. 3.**
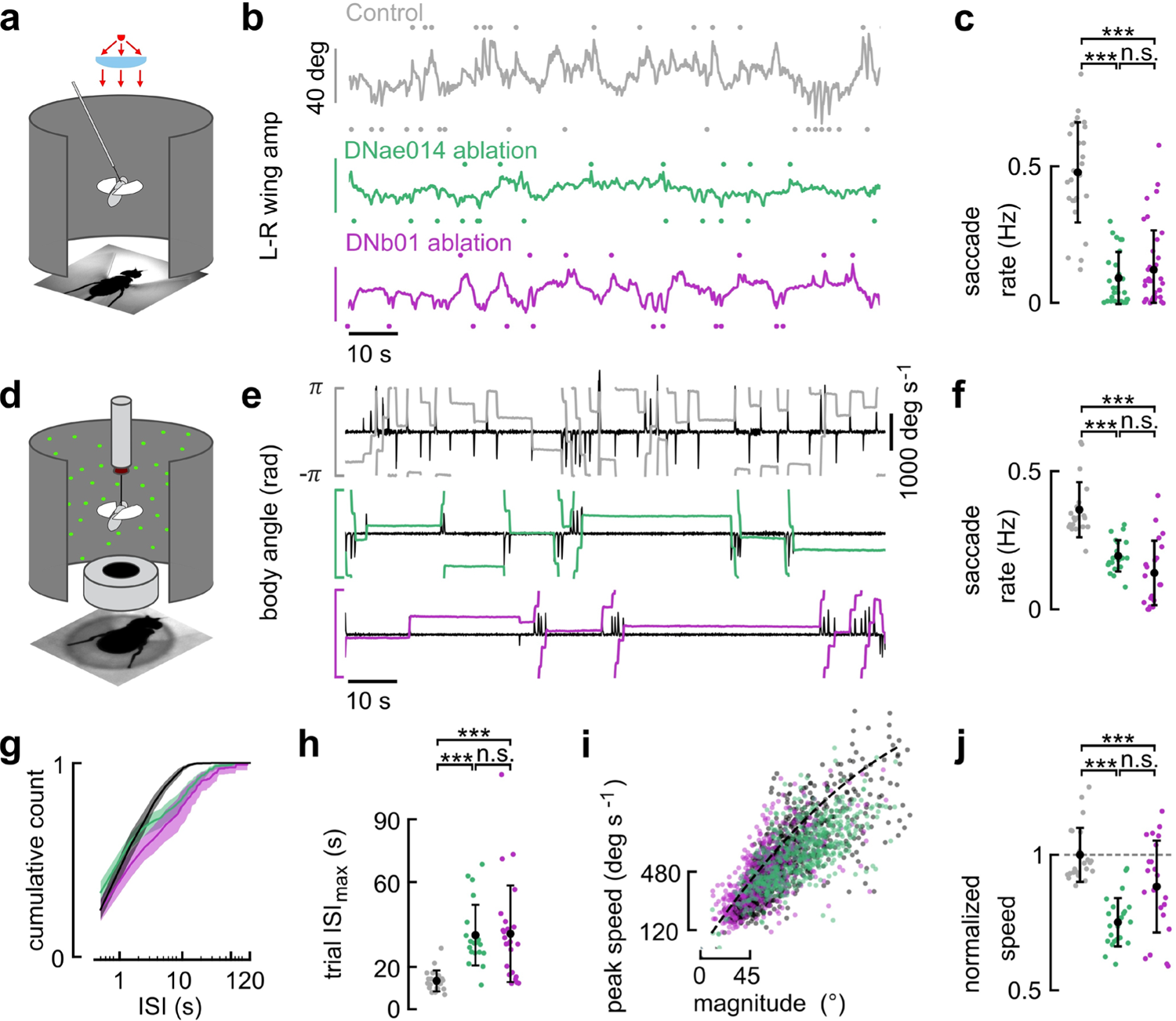
Ablation of either DNae014 or DNb01 alters saccade dynamics. **a**, Riged-tether flight arena with IR backlighting to track wingstroke amplitudes. **b**, Transient changes in L-R wingstroke amplitude indicate fictive saccades. Example control fly (top, grey), DNae014-ablated fly (middle, green), and DNb01-ablated fly (bottom, magenta) **c**, Individual saccade rates in the rigid tethered arena (n = 26, 25, 36, respectively). **d**, Magnotether flight arena with IR backlighting to track body angle and a static visual pattern. **e**, Example traces with saccades apparent as rapid changes in body angle and angular velocity transients (black traces). **f**-**j**, Color codes as in **b**. n = 22, 23, 22 flies, respectively. **f**, Saccade rates in the magnotether arena, plotted as mean of individual fly means and boot-strapped 95% CI. **g**, Cumulative histogram of inter-saccade intervals. **h**, Maximum inter-saccade interval (ISI_max_) for each 2-minute trial. **i**, Relationship between peak saccade speed and saccade magnitude. Dashed line: Orthogonal-distance regression fit of a 2^nd^ order polynomial to pooled data of control flies. **j**, Peak saccade speed normalized to fit in **i,** plotted as mean of individual fly means and boot-strapped 95% CI. Statistical differences were assessed using Kruskal-Wallis tests with Tukey’s HSD method for post-hoc comparisons (**c**, **f**, **h**, **j**; *** = p<0.001; n.s. = not significant).

To gain further insight into the effect of cell ablation on saccade generation, we repeated the experiments using a magnotether arena, in which both rapid turns and epochs of straight flight are easier to observe and classify (Bender and Dickinson, 2006) (Fig. 3d, e). Ablating the DNs reduced the saccade rate in the magnotether from 0.37 ± 0.10 Hz in empty vector controls to 0.20 ± 0.06 Hz for DNae014-ablated flies (p<0.001) and 0.19 ± 0.10 Hz for DNb01-ablated flies (p<0.001; Fig. 3e, f). In addition, genetic ablation of either cell type resulted in a qualitatively different temporal pattern of flight behavior distinguished by long sequences of straight flight interspersed with rapid bursts of saccades, typically in the same direction (Fig. 3e, g). We quantified this influence of cell ablation on saccade timing by determining the maximum inter-saccade interval (ISI_max_) for each two-minute trial of each fly. The ISI_max_ for control flies was 13.2 ± 4.9 s, compared to 33.1 ± 15.2 s for DNae014-ablated flies (p<0.0001) and 33.1 ± 22.1 s for DNb01-ablated flies (p<0.0001, Fig. 3h). As observed previously (Bender and Dickinson, 2006), the speed and magnitude of saccades in the magnotether were highly correlated, with faster saccades being larger (Fig. 3i). The saccades after DNae014 ablation were slower than controls, as were those following DNb01 ablation (p<0.001 and p=0.009, respectively; Fig. 3j). The altered saccade dynamics and temporal distribution after ablation support the hypothesis that each DN can produce saccades independently of the other, but with different dynamics than those of control flies. The results support a working hypothesis that the two DNs play complementary roles by activating different components of the motor circuit in the VNC responsible for generating saccades. Together, functional imaging, unilateral activation, and ablation experiments suggest that two pairs of descending interneurons, DNae014 and DNb01, function together as saccade-generating couplets (SGCs) to execute commands for spontaneous turns during flight.

To further examine the underlying circuitry in the brain, we identified the two pairs of DNs in two available connectomes. Comparing confocal image stacks of GFP expression in DNae014 to the FlyWire and hemibrain connectomes (Scheffer et al., 2020; Dorkenwald et al., 2023; Schlegel et al., 2023) revealed at least four DN pairs with similar morphology, all within the DNa class as defined by Namiki and coworkers (Namiki et al., 2018). Of these, only one (DNae014) appeared to be a precise match with the cell originally described as AX by Schnell and coworkers (Schnell et al., 2017) (Gregory Jefferis, pers. comm.), which is why we chose to adopt the new nomenclature (Dorkenwald et al., 2022; Dorkenwald et al., 2023; Schlegel et al., 2023) (Eichler, Brooks, Stürner *et al*., in prep.). DNae014 appears to be identical to the PS017_R neuron in the hemibrain connectome. Our confocal microscopy images of DNb01 matched only one DN in the two brain connectomes, which was already annotated as DNb01 in both the hemibrain and FlyWire datasets. Based on the results of our physiological and behavioral experiments (Figs. 1-3), we propose that the DNae014 and DNb01 cells with somata on the left side of the brain form a functional couplet that generates leftward saccades, with the right DNae014 and DNb01 cells forming a couplet that generates rightward saccades. Neurotransmitter predictions based on machine learning (Buhmann et al., 2021) suggest that DNae014 is cholinergic and DNb01 is glutamatergic (Extended Data Table 1). These neurotransmitter types indicate that the two members of each couplet collectively carry excitatory and inhibitory commands to opposite sides of the wing neuropil in the VNC. Recent evidence from two new connectomes of the adult VNC suggest, but do not confirm, that these two cells act on steering motor neurons both directly and indirectly via a large population of premotor interneurons (Cheong et al., 2023; Lesser et al., 2023). Because both inhibitory and excitatory premotor neurons exist, the valence of a DN’s neurotransmitter is not necessarily a clear predictor of its action on steering muscle activity during saccades. Within the brain, the DNb01 cells synapse directly onto contralateral DNae014 cells. This is a simple reciprocal inhibitory motif consistent with a network responsible for binary turning to either the left or the right (Extended Data Table 2), analogous with the arrangement of Mauthner cells in fish (Korn and Faber, 2005).

Motivated by our hypothesis that each unilateral set of DNae014 and DNb01 cells functions as a saccade-generating couplet (SGC), we performed a connectivity search for neurons that provide direct inputs to both cells in a SGC (Fig. 4). Basing our analysis on the lowest synapse number threshold that maintained a bilaterally symmetric connectivity pattern (see Methods for details, Extended Data Fig. 4a), we identified 8 neuron types that provide direct input to the SGCs, (a diverse population we call SGCIs). Five of these SGCIs are pairs of neurons with unique morphology and three constitute small sets of cell types with nearly identical morphology and connectivity (Extended Data Fig. 4b). In addition to making direct synapses with the two members of each SGC, several SGCIs send a collateral that directly connects with other SGCI members on the ipsilateral or contralateral side (Fig. 4). We further identified a class of intermediary neurons that form strong connections between members of contralateral SGCIs. Predictions of transmitter type based on machine learning suggest that the population of SGCIs and intermediary cells includes both excitatory (acetylcholine) and inhibitory (glutamate or GABA) neurons (Fig. 4; Extended Data Fig. 5). We also ranked the top ten additional neuron types that form presynaptic connections with members of this putative saccade network. These inputs include neurons that likely carry sensory information from brain regions including the lobula (Lo), posterior lateral protocerebrum (PLP), accessory optic tubercle (AOTu), as well as the LAL and central complex (CX) (Fig. 4i; Extended Data Fig. 6). A recent study also identified neuron types (PFL2 and PFL3) that carry information from the CX to DNs in the LAL (Hulse et al., 2021). In addition, DNb01 and several SGCIs make recurrent inhibitory connections to network members on the contralateral and ipsilateral sides (Extended Data Fig. 5a). These anatomical pathways suggest that external sensory information and internal processing likely influence the dynamics of the network that generates spontaneous saccades within the LAL.

**Fig. 4.**
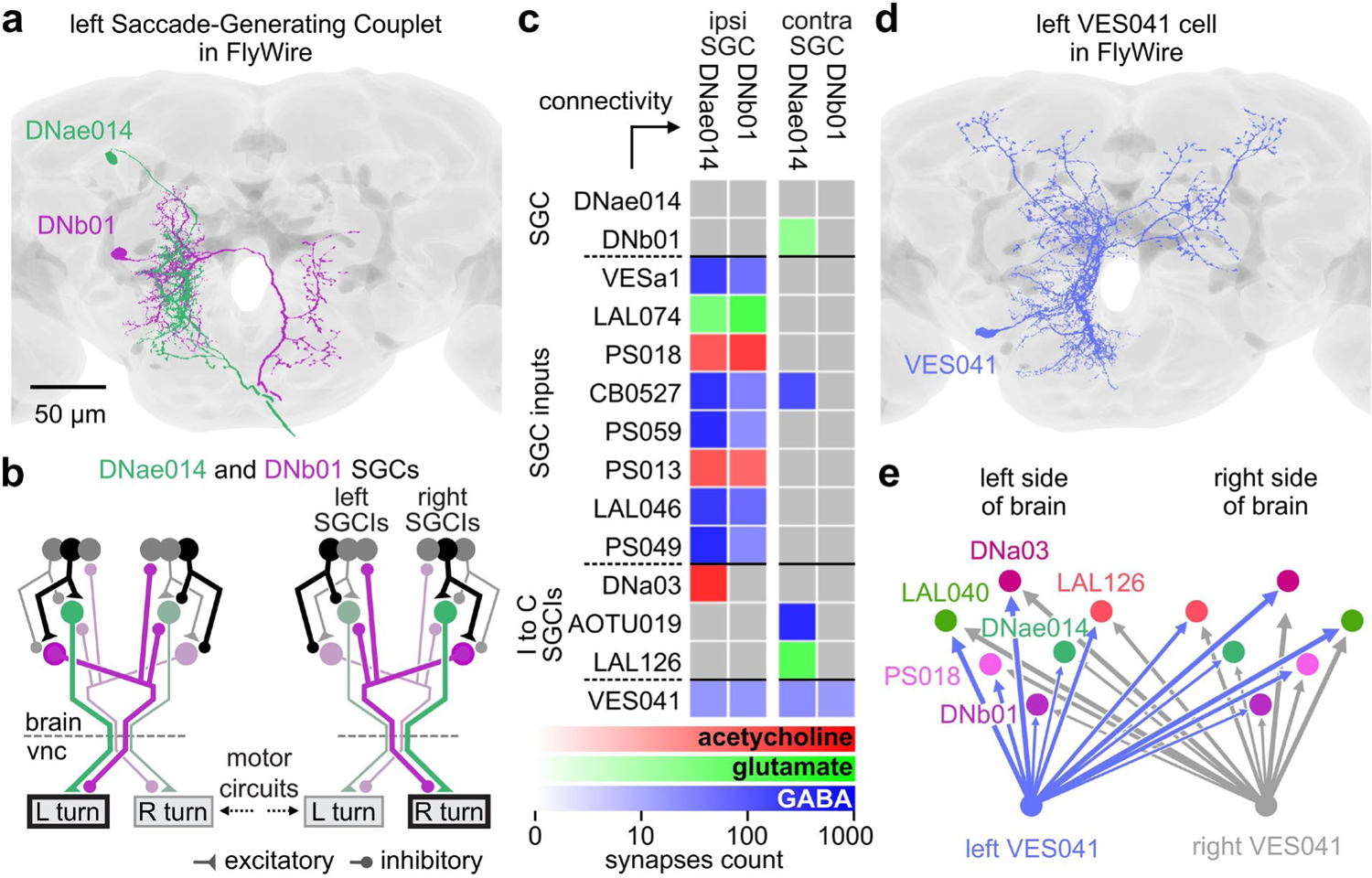
VES041 neurons innervate all DNae014 and DNb01 cells, along with several of their upstream targets. **a**, Left saccade-generating couplet (SGC) consisting of DNae014 (green) and DNb01 (magenta) cells (from FlyWire). **b**, Schematized model for network generating left and right saccades. Bright colors or black indicate active neurons; pale colors or gray indicate inactive neurons. **c**, Connectivity of putative members of the network that regulates directed saccades (rows) to SGC neurons DNae014 and DNb01 (columns; arrow indicates direction of information flow). Our analysis identified neurons that form input synapses to both SGC cells (SGCIs), neurons that form connections between ipsilateral and contralateral SGCIs (I to C SGCIs), and a neuron that connects to all four SGC cells (VES041). Connectivity data are averaged assuming symmetrical arrangement of left and right network members. **d**, Morphology of the left VES041 cell in FlyWire. **e**, Connections of VES041 cells (blue for left; grey for right) to both SGCs and bilateral pairs of SGCIs and intermediary neuron types. Colors match representations in Extended Data Fig. 5c-j. Line thickness is proportional to log of synapse count.

One cell, VES041, stood out in our analysis due to its unique property of providing input to the DNae014 and DNb01 cells of both SGCs as well as to several of their prominent upstream targets (Fig. 4c, e; Extended Data Fig. 5). The neurotransmitter for VES041 is predicted to be GABA (Extended Data Table 2). Thus, each of the paired VES041 cells could potentially inhibit all four cells that constitute the two SGCs. Because of this connectivity, we hypothesized that prolonged activation of VES041 would strongly suppress saccade rate. To test this, we generated two split-GAL4 lines that target VES041 neurons and used them to drive CsChrimson (see Methods; Fig. 5A). Presentation of 617 nm light strongly reduced spontaneous saccades in flies flying in a magnotether apparatus in both driver lines (Fig. 5b-f). These results are consistent with independent experiments conducted on rigidly tethered flies in which we classified saccades based on changes in wingstroke amplitude (Extended Data Fig. 7). These experiments thus support our hypothesis that VES041 is a potent inhibitor of spontaneous saccades. The VES041 neuron receives its strongest input from the vest (VES), GNG, LAL, SPS, flange (FLA), inferior bridge (IB), and saddle (SAD) (Fig. 5; Extended Data Fig. 8), which suggests that this cell integrates both visual and olfactory information that are known to influence saccade rate.

**Fig. 5.**
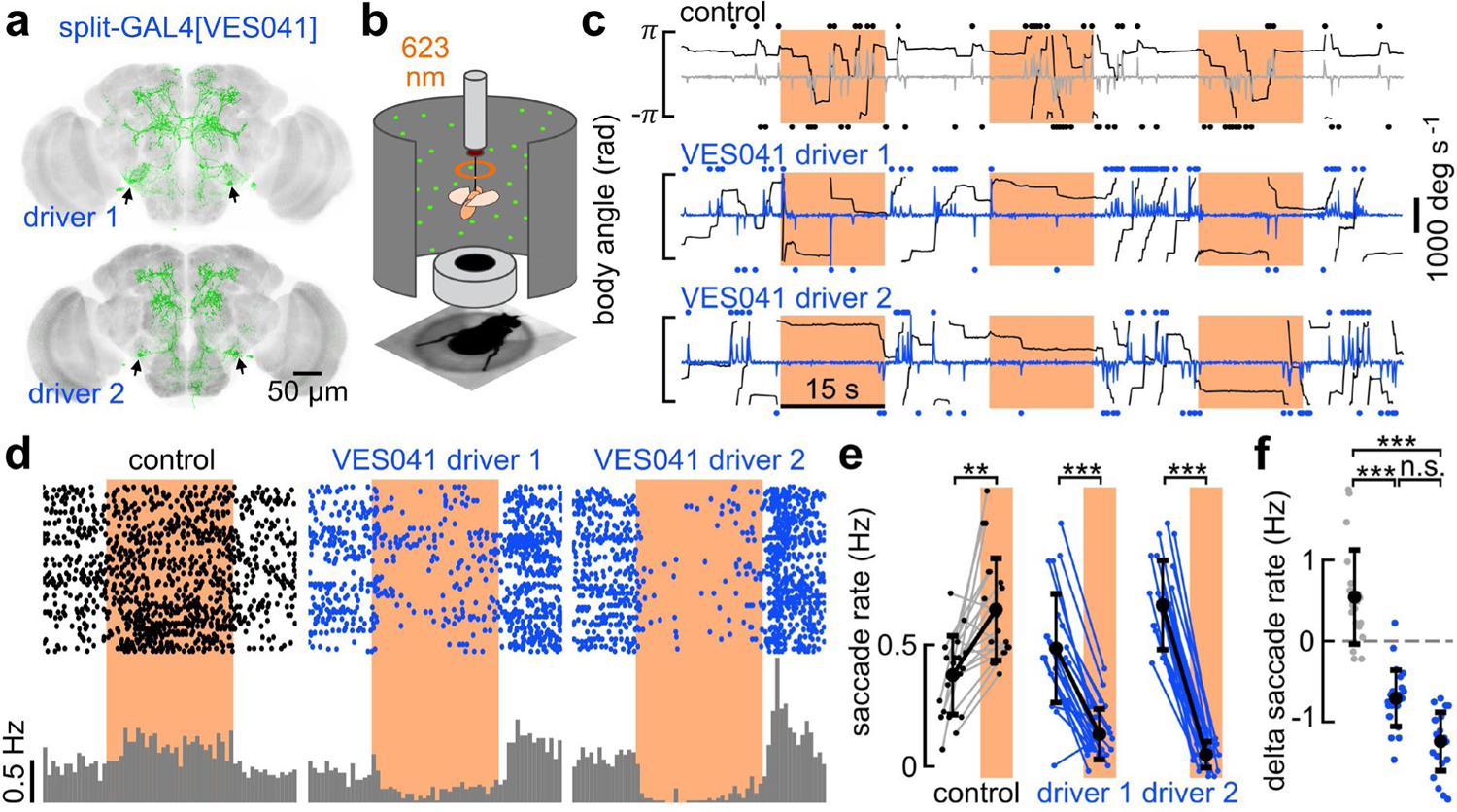
Activation of GABAergic VES041 neurons suppresses spontaneous saccades. **a**, Split-GAL4 lines constructed to target VES041 (green, GFP). **b**, Optogenetic activation of VES041 in a magnotether arena using a 623 nm LED ring light (orange annulus in **b**, orange patches in **c**). **c**, Top: empty vector control fly, with right and left saccades (dots above and below the panel) identified using angular velocity of body orientation (grey trace). Middle and bottom: activation of VES041 drivers 1 and 2 (angular velocity in blue). **d**, Raster plots represent spontaneous saccades (black dots), during exposure to 623 nm light (orange patch). Each set of three rows are the presentations for an individual fly. Left: empty vector split-GAL4 line driving CsChrimson; middle and right: VES041 split-GAL4 lines 1 and 2 driving CsChrimson. Gray histograms indicate mean saccade rate across populations. **e**, Individual saccade rates during light-off and light-on epochs for control and the two VES041 drivers; Each line represents an individual fly (n = 20, 24, and 23 flies). **f**, Individual differences in saccade rate from the light-off to light-on condition for the same groups as in **e.** (** = p<0.01; *** = p<0.001; n.s. = not significant, Kruskal-Wallis test followed by post-hoc comparisons with Tukey’s HSD method).

## Discussion

We investigated the role of two pairs of DNs in the generation of body saccades in flying flies. Our results suggest a model in which two neurons, DNae014 and DNb01, form two saccade-generating couplets (SGCs). Although, we could not record from DNae014 and DNb01 simultaneously, every rapid change in wing motion accompanied a transient change the activity of either cell when recorded separately. This one-to-one correspondence supports the hypothesis that DNae014 and DNb01 function as single command units that carry both excitatory and inhibitory inputs to the motor circuits in the VNC that execute flight saccades. In the brain, we used the DNae014 and DNb01 cells to identify a mosaic of excitatory and inhibitory neurons that innervate neuropils associated with steering and multi-modal sensory integration. One pair of large inhibitory interneurons, VES041, emerged as a likely candidate to suppress the execution of spontaneous turns because each cell makes strong inhibitory connections with all members of both the left and right SGCs. Optogenetic activation of VES041 strongly suppresses saccades, which suggests that it may indeed play a regulatory role in natural behaviors such as long-distance dispersal or upwind surges in response to an attractive odor. Although we have not yet succeeded in performing a viable double ablation experiment (i.e. silence or ablate both DNae014 and DNb01 in the same individuals) the results of VES041 activation serve as a rough proxy for simultaneous silencing of both DNae014 and DNb01 (Fig. 5, Extended Data Fig. 7).

While the high fidelity of the activity of both cells to the changes in wing motion is itself compelling (Fig. 1), this phenomenon might result from strong ascending proprioceptive feedback or efferent copy. However, brief activation of either single DNae014 or single DNb01 cells reliably evoke syndirectional turns (Fig. 2). Further, ablating either cell type results in unusually long sequences of straight flight interspersed by brief bursts of small amplitude saccades (Fig. 3). The induction of long inter-saccade intervals provides intriguing evidence that these cells contribute directly to temporal patterning of the spontaneous saccades and do not simply relay upstream commands (Figs 1, 2, and 3, Extended Data Fig. 2). However, ablating either cell alone does not abolish all spontaneous turns, even though the persisting saccades are slower in speed. This partial loss-of-function effect might indicate that the motor commands resulting from either DNae014 or DNb01 activate sub-components of the complete motor program that are alone sufficient to generate partial, slower saccades (Fig. 3j), or alternatively, that the fly’s nervous system contains as-of-yet unidentified parallel pathways for generating spontaneous turns. These alternative hypotheses are not mutually exclusive and, because our ability to cluster saccades according to type based on behavioral measurements is limited, not currently distinguishable.

In summary, our results suggest that DNae014 and DNb01 form two saccade-generating couplets that are inhibited by VES041. DNae014 and DNb01 cells are probably members of a quite ancient network that is centered in the paired LALs of the brain and widespread among insects and other arthropods (Steinbeck et al., 2020; Strausfeld and Hirth, 2013). In particular, comparative anatomical findings suggest these two pairs of DNs are homologous to ‘flip-flop’ neurons responsible for the rapid turns exhibited during plume tracking behavior in the silkmoth, *Bombyx mori* (Kanzaki, 1998; Olberg, 1983). Future experiments are needed to identify the specific sensory cues and their anatomical pathways that influence VES041 activity, and thus confirm its role in regulating this ecologically important behavioral state transition in flying insects.

## Acknowledgements

We thank members of our lab, including Ainul Huda, Will Dickson, Johan Melis, Francesca Ponce, and Matthew Clark. Additionally, we are grateful to Shigehiro Namiki, Sasha Rayshubskiy, Rachel Wilson, and Anna Ahn for helpful contributions and discussions, and to Sigehiro Namiki, Wyatt Korff, and Gwyneth Card for sharing the driver lines used in the functional imaging screen. We thank the Princeton FlyWire team and members of the Allen Institute for Brain Science for development and maintenance of FlyWire (supported by BRAIN Initiative grants MH117815 and NS126935 to Murthy and Seung). We also acknowledge members of the Princeton FlyWire team and the FlyWire consortium for neuron proofreading and annotation, specifically J. Dolorosa, D. Sapkal, S. Fang, and Z. Vohra from the Murthy, Seung, and Jefferis labs. This research was supported by NIH grants U19NS104655 and 1R21NS106471-01A1.

## Author contributions

(Following CRediT taxonomy): conceptualization: I.G.R., M.H.D.; methodology: I.G.R., J.J.O. M.H.D.; software: I.G.R.; validation: I.G.R, J.J.O.; formal analysis: I.G.R.; investigation: I.G.R, J.J.O.; resources: I.G.R., J.J.O.; data curation: I.G.R.; writing original draft: I.G.R., M.H.D.; writing review & editing: I.G.R., M.H.D.; visualization: I.G.R and J.J.O.; funding acquisition: M.H.D.; supervision: M.H.D.; project administration: M.H.D.

## Competing interests

The authors have no competing interests to declare.

## Methods

### Experimental model details

All experiments were conducted using 2-to-5-day-old adult female *Drosophila melanogaster.* To specifically target DNae014 cells, we constructed a split-GAL4 line using the hemidrivers VT025718-GAL4.AD to drive the activation domain, and R56G08-GAL4.DBD to drive the DNA-binding domain, resulting in split-GAL4[VT025718.AD; R56G08.DBD] (Dionne et al., 2018; Namiki et al., 2018; Pfeiffer et al., 2010; Tirian and Dickson, 2017). Similarly, we created two split-GAL4 drivers that target VES041 cells, split-GAL4[R64G09.AD; VT044519.DBD] and split-GAL4[R64G09.AD; R31E10.DBD]. Candidate hemi-drivers followed from screening databases of standardized MCFO confocal scans of expression patterns of GAL4 drivers (Clements et al., 2022; Nern et al., 2015). All hemidrivers were obtained from the Bloomington Drosophila Stock Center (Cook et al., 2010). For simplicity, we refer to these specific drivers by their primary targeted neurons, while acknowledging the potential presence of off-target neurons (Extended Data Videos 1-8). We used the previously generated split-GAL4 driver SS02383 to target DNb01 cells (Namiki et al., 2018). For functional imaging experiments, we crossed males of either the DNae014 split driver or the DNb01 split driver to w+;UAS-tdTomato;UAS-GCaMP7f female flies (Namiki et al., 2022). For stochastic optogenetic activation experiments, we crossed males from the respective split-GAL4 lines combined with nSyb-PhiC31 (X) with virgin females carrying the intermediate or sparse variants of 20XUAS-SPARC-Syn21-CsChrimson::tdTomato-3.1 (Isaacman-Beck et al., 2020). For VES041 optogenetic activation experiments, we crossed males from the two VES041 split drivers with females carrying 20XUAS-CsChrimson-mVenus on the third chromosome. For temporal silencing experiments, we drove expression of the light-gated GtACR1 channel by crossing males of the DNae014 split driver with females carrying 20XUAS-GtACR1-EYFP on the third chromosome (Govorunova et al., 2015). For genetic ablation experiments, we drove expression of the pre-apoptotic genes *reaper* and *hid* by using female offspring from crosses between males from each of the DNae014 and DNb01 split drivers with females carrying UAS-rpr,hid on the first chromosome (Goyal et al., 2000). We reared the progeny of experimental crosses on standard cornmeal fly food, supplemented with additional yeast. For optogenetic activation experiments, larvae were reared in food supplemented with 0.2 mM all trans-Retinal (ATR) (Sigma-Aldrich). For the first two days following eclosion, adult flies were transferred to bottles with standard cornmeal fly food supplemented with 0.4 mM ATR.

### Quantification and statistical analyses

All experiments were analyzed with custom software written in Python. Sample sizes refer to the number of individuals tested. Variance across individuals was assessed by either standard deviation of individual means or boot-strapped, 95% confidence intervals (CI) for the mean of the individual means (Figs. 3c, f, h, and j, 5e, and Extended Data Fig. 3d, and Figs. 1m, Fig. 2c, g-i, 4i, and Extended Data Figs. 2b-d).

## Methods details

### Anatomy

The central nervous system of 3-to-5-day-old female adult progeny were dissected at room temperature (∼20°C) in 1X PBS (Sigma) and fixed for 25 min in 4% paraformaldehyde diluted in 1X PBS, and washed 3 times for 15 min in 1X PBS and subsequently 5 times for 15 min in 0.3% PBS-T (1X PBS with 0.3% Triton-X). Samples were then blocked in 7.5% normal goat serum diluted in 0.3% PBS-T for at least 1 hour. The samples were incubated with primary antibodies (mouse nc82 supernatant at 1:30, rabbit polyclonal anti-DsRED at 1:2000 or chicken anti-GFP 1:1000, diluted in blocking solution) for 4 hours at 20°C and 3 days at 4°C, followed by incubation with secondary antibodies (Cy5 goat anti-mouse at 1:300, and Alexa Fluor 546 goat anti-rabbit at 1:1000 or Alexa Fluor 488 goat anti-chicken at 1:1000, diluted in blocking solution) for 4 hours at 20°C and 3 days at 4°C. Samples were imaged using a confocal microscope (Zeiss LSM 880) with a 40X water objective. Optical sections were taken at 1-μm intervals with resolutions of 512X512 or 1024X1024 pixels. The samples were mounted on microscope slides with spacers using Vectashield Plus (VectorLabs).

### Functional imaging

Tethered, flying flies were imaged using a galvanometric scan mirror-based two-photon microscope (Thorlabs) equipped with a Nikon CFI apochromatic, near-infrared objective water-immersion lens (40x mag., 0.8 N.A., 3.5 mm W.D.), as previously described (Namiki et al., 2022; Schnell et al., 2017). GCaMP7f and tdTomato fluorescence signals were imaged in the arbors of the left and right DNae014 or DNb01 cells in the SPS neuropil in the brain. Imaging parameters varied depending on the preparation; we either acquired 72 x 36 µm images with 128 x 64 pixel resolution at 13.1 Hz, or in a few cases, 36 x 36 µm images with 96 x 96 pixel resolution at 10.6 Hz (Extended Data Fig. 1). To correct for brain motion in the x-y plane, we registered both channels for each frame using a cross-correlation between each tdTomato image and the trial-averaged image (Guizar-Sicairos et al., 2008). A field of view (FOV) centered along the bilateral axis was selected based on tdTomato expression. Regions of interest (ROIs) to capture signals of the left and right cells were automatically determined based on GCaMP7f fluorescence pixel variance and covariance with turning behavior, refining signal-to-background ratios and reducing cross talk between contralateral DNb01 cell clusters (see below for details). To account for motion in the z-axis, we normalized GCaMP7f fluorescence to tdTomato fluorescence. To find the baseline fluorescence (F_0_), we subtracted the background fluorescence, defined as the 20% dimmest pixels of the entire FOV, from the mean of the lowest 5% fluorescence values in a trial. To standardize neuronal activity, we normalized baseline-subtracted fluorescence, F_t_, to the maximum observed for each individual fly in each ROI: ΔF/F = (F_t_ – F_0_) / (F_t_ – F_0_) _max_. Neural activity during spontaneous flight turns was recorded in 90 s trials in darkness (i.e. the microscope was enclosed in a light-tight enclosure and the visual arena was turned off). To measure responses to looming, files were presented with an expanding stimulus created using a 96 x 32 pixel arena that covered 216° of azimuth and ∼74° of elevation(Reiser and Dickinson, 2008). Looming stimuli simulated a 0.15 m diameter disk approaching at 1.5 ms^-1^ from a starting position of 1.5 m away from the fly; each trial ended when the stimulus subtended an angle of 60°. Looming stimuli were presented every 6 seconds from one of five directions in pseudo-random order. To reduce light pollution, we shifted the spectral peak of the visual stimuli from 470 nm to 450 nm by covering the arena panels with transmission filters (one sheet of Roscolux no. 59 Indigo, two sheets of no. 39 Skelton Exotic Sangria, and two sheets of no. 4390 Cyan). We tracked left and right wingstroke amplitudes with a machine vision system, Kinefly, as described before (Suver et al., 2016).

### Functional Imaging, regions of interest

We refined the imaging ROIs for functional imaging via an automated process that used the covariance between wingstroke amplitude and pixel variance in GCaMP7f fluorescence, which we confirmed via direct inspection of cell anatomy (Extended Data Fig. 1). Initially, we manually assigned ROIs to capture activity of left and right cells based on visual assessment of tdTomato and GCaMP7f intensities. For the tdTomato signal, we determined the maximum intensity projection in the red channel across all images for each flight trial. For the GCaMP7f signal, we determined the standard deviation for every pixel across each flight trial. We then set a threshold for GCaMP7f intensity within the interrogation areas to identify active pixels and created a binary mask for the left and right cells (Extended Data Fig. 1b). These manually constructed ROIs were used to calculate normalized fluorescence signals (ΔF/F) as described earlier. Relationships between neural activity and behavior were determined using ΔF/F based on manual ROIs through cross-correlation with wingstroke amplitude (Extended Data Fig. 1c). The ΔF/F signal in the manually assigned ROI correlated positively with contralateral wingstroke amplitude, and contralateral minus ipsilateral wingstroke amplitude, for both DNae014 and DNb01. To refined single-cell ROIs, we constructed pixel correlograms, based on time-varying fluorescence within a small set of pixels assessed to belong to a single cell, or, for automatic assignment of DNae014 and DNb01 morphology, we cross-correlated the contralateral minus ipsilateral wingstroke amplitude with all pixel fluorescence values from the GCaMP7f channel in a given flight trial. Top ranked pixels from the resulting correlograms were thresholded and binarized to create ROIs for the left and right cells. These ROIs were visually validated against cellular anatomy (Extended Data Fig. 1e). Automatic ROIs were used to obtain single cell GCaMP7f ΔF/F separately for DNae014 and DNb01 trials, following the same procedures described earlier.

### Saccade detection

Fictive saccades were detected using a threshold-based classifier (Schnell et al., 2017) operating on the temporal derivative of the bilateral difference in wingstroke amplitude. We used a threshold of twice the measured standard deviation of the derivative for both the functional imaging setup and the rigid tether arena. To avoid tracking errors, we excluded derivative values exceeding 420 ° s^-1^ and cases in which stroke amplitude fell below 60°. To avoid false positives, caused by rapid changes in wingstroke angles at the end of saccades, we implemented a 0.25 s refractory period following each saccade as in earlier studies (Kim et al., 2015). Wingstroke amplitude signals were smoothed using a Kalman filter that approximated a 4^th^-order Butterworth with a low-pass cut-off at 6 Hz. Saccades were classified similarly for magnotether data, using the time derivative of the body angle and a threshold of 1.5 standard deviations (Fig. 3e). Peak saccade speeds in the magnotether were detected by identifying extrema in body angle velocity after applying a Savitsky-Golay filter (< 9 Hz). We normalized peak saccade speed to expected peak speed for a given saccade magnitude, based on the measured relationship between saccade speed and saccade magnitude for control flies. Specifically, we normalized all saccade data to the orthogonal-distance regression (ODR) (Boggs et al., 1988) between peak speed and magnitude for all saccades measured in empty-vector control flies (Fig. 3i). We used an 2^nd^-order polynomial in the ODR because it matched the curvilinear relationship better than a simple line.

### Optogenetic activation experiments

Tethered flies were positioned for imaging with a digital camera and backlit with infrared light. Prior to CsChrimson activation, we presented flies with an 8 s visual starfield pattern oscillating sinusoidally about the vertical axis through two full cycles of 216° in amplitude. Wingstroke amplitudes were tracked using Kinefly software (Suver et al., 2016). For optogenetic activation in the rigid tether setup, we guided a fiber optic from a 617 nm LED source (M617F2, Thorlabs) through a microscope objective, adjusted to create a 0.5 mm spot on the posterior side of the head (Fig. 2d and Extended Data Fig. 7), with a power density of ∼0.2 mW mm^-2^. For optogenetic activation in the magnotether, we delivered 623 nm light via a custom-built LED ring (Fig. 5b). For unilateral activation experiments of either DNae014 or DNb01 (Fig. 2d-i) we elicited 6 responses to a 0.25 s light pulse at 10 s intervals. Bilateral CsChrimson activation experiments of VES041 involved three 15 s periods of excitation surrounded by 15 s periods of no excitation. If a fly stopped flying, we collected additional trials. For each fly, the three first presentations with continuous flight were included for further analysis.

### Unilateral genetic targeting of CsChrimson

To stochastically target the optogenetic effector CsChrimson tagged with tdTomato within the split-GAL4 driver lines (for either DNae014 or DNb01), we employed Sparse Predictive Activity through Recombinase Competition (SPARC) (Isaacman-Beck et al., 2020). We crossed parental lines containing pan-neuronal phi-C31 and the split-GAL4 of interest with the appropriate SPARC line (20XUAS-SPARC2-I-Syn21-CsChrimson::tdTomato-3.1 for intermediate labeling or (20XUAS-SPARC2-D-Syn21-CsChrimson::tdTomato-3.1 for dense labeling). The resulting flies expressed CsChrimson and a fluorescent marker, tdTomato, in either the left cell, the right cell, both cells, or no cells. Expression patterns were manually scored using fluorescence microscopy after behavioral measurements, and a subset of flies underwent immunostaining to obtain representative confocal microscopy images (Fig. 2e). All experiments were performed in a blinded fashion, with expression patterns naïvely scored behavioral data collection. These experiments were performed on intact, rigidly tethered flies.

### Connectomics

We identified the DNae014 and DNb01 cells in publicly available connectomes (Scheffer et al., 2020; Dorkenwald et al., 2023; Schlegel et al., 2023). Comparison to fly brain connectomes indicated at least five DNs in the subclass of DNa neurons (Namiki et al., 2018) that roughly resembled DNae014, but one neuron matched the morphology of the cell in our split-GAL4 line most closely (Gregory Jefferis and Marta Costa, pers. comm.). The hemibrain dataset contains two neurons labeled DNb01, with one matching the expression pattern of the cell we refer to a DNb01 based on the description of Namiki at coworkers (Namiki et al., 2018), which is under control of the split-GAL4 driver we used in our experiments, SS02383. We conducted a connectivity analysis based on both the FlyWire connectome based on the FAFB EM dataset, and the hemibrain connectome datasets (Dorkenwald et al., 2022; Dorkenwald et al., 2023; Scheffer et al., 2020; Schlegel et al., 2023; Zheng et al., 2018) (see flywire.ai). Neurons in FlyWire were considered of the same type following labeling in the hemibrain and/or due to similarity in both anatomy and connectivity (see Extended Data Fig. 4a). We identified saccade-generating couplet input neuron types (SGCIs) that innervated both unilateral DNae014 and DNb01 cells with combined connection strengths above a threshold where the product of synapse counts onto DNae014 and DNb01 cells exceeded 75^2^. This threshold was objectively determined as the lowest threshold at which the set of SGCIs to the left SGC were identical to the symmetrical set of SGCIs to the right SGC (Extended Data Fig. 4b, c). Putative saccade network neurons were labeled according to the FAFB or hemibrain connectomes (Extended Data Table 3). To assess interconnectivity between identified neurons, we included direct connections that contained over 37 synapses (Extended Data Fig. 5, Fig. 4), as well as indirect connections between left and right SGCIs that contained over 75 synapses. For these interneurons connecting left and right SGCIs, we applied a higher threshold to exclude more likely false positive connections due to high local innervation density. Neurotransmitter types were predicted using a machine-learning synapse classification algorithm based on confidence scores and synapse cleft width (Buhmann et al., 2021; Eckstein et al., 2020; Heinrich et al., 2018).

## Extended Data figures

**Extended Data Fig. 1.**
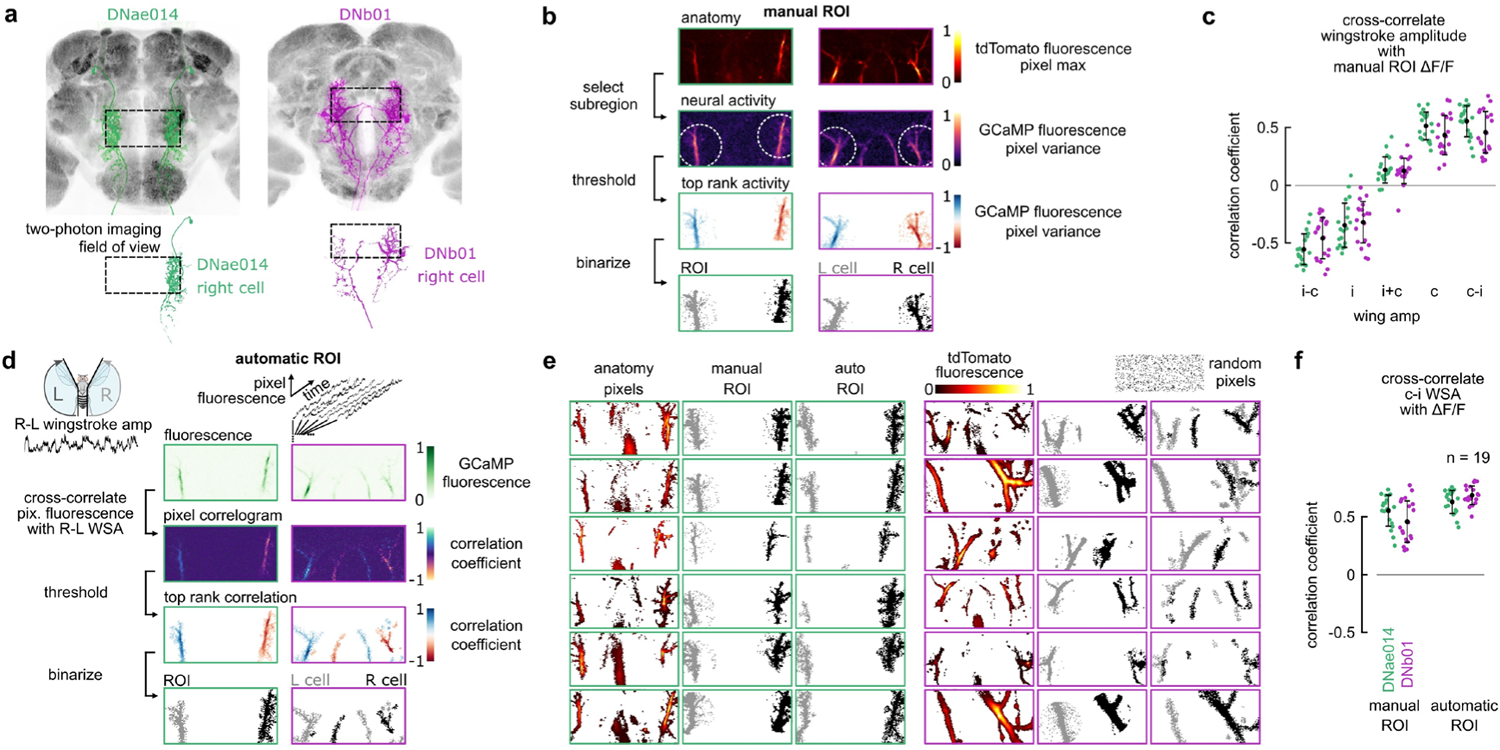
Automatic ROI determination using neural activity variance and covariance with wingstroke amplitude signals. We refined ROIs for functional imaging, to improve single cell fluorescence signals. This automated procedure uses covariance between wingstroke amplitude (WSA) and pixel variance in GCaMP7f fluorescence, and is verified via inspection of cell anatomy. **a**, Approximate 2-photon functional imaging Field of View (FOV) shown by dashed box encompassing bilateral anatomical expression of DNae014 (left, green) and DNb01 (right, magenta). Insets below with example single-cell expression patterns with the FOV. **b**, Manual assignment of Regions of Interest (ROIs) for DNae014 (green outlines) and DNb01 (magenta outlines). Interrogation areas (white dashed circles), based on tdTomato fluorescence maximum intensity projection (top) and GCaMP7f fluorescence variance projection (2nd row), following tdTomato-registration (see Methods). Top ranked GCaMP7f fluorescence variance within interrogation areas binarized to create manual ROIs for left and right cells. **c**, Normalized coefficients for cross-correlations between GCaMP7f signals from manually assigned ROIs and WSAs for DNae014 (green, n=19) and DNb01 (magenta n=19). Abbreviations for WSA signals relative to the cell body location: i-c, ipsilateral-contralateral; i, ipsilateral, i+c, ipsilateral+contralateral; c, contralateral, c-i, contralateral-ipsilateral. **d**, Automatic ROI determination for example DNae014 (green outlines) and DNb01 (magenta outlines). Pixel-wise cross-correlations between GCaMP7f fluorescence vectors and bilateral WSA difference (R-L) used to automatically determine ROIs based on positive and negative correlations. **e**, Comparison of top ranked fluorescence pixels in tdTomato mean-projection images with manual ROIs (middle column), automatic ROIs (right columns), and a random pixel pattern (right top). **f**, Cross-correlation coefficients between contra-ipsilateral WSA and ΔF/F based on manual ROIs (left) and automatic ROIs (right), for DNae014 (green) and DNb01 (magenta). **e**, **f,** ROI refinement only marginally improves ROIs in matching the morphology of DNae014 and the association between behavior and DNae014 activity. However, for DNb01, with intertwined morphologies of each cell innervating both side of the brain, automated ROI detection more accurately reflects single cell morphology and increases the association between behavior and DNb01 activity.

**Extended Data Fig. 2.**
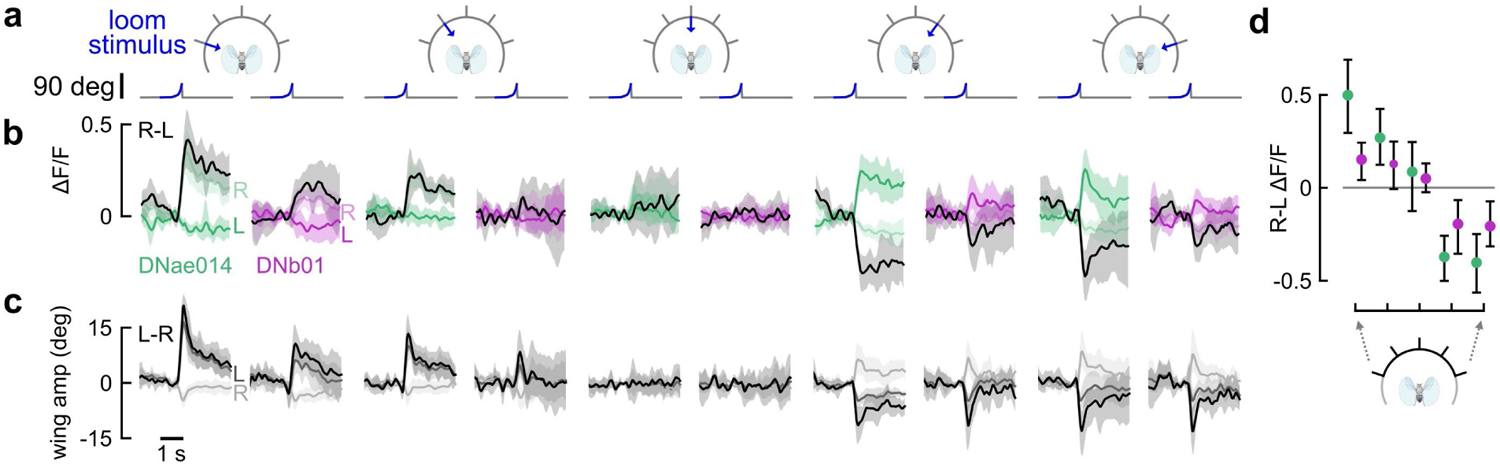
DNae014 and DNb01 are active during saccades elicited by visual loom. **a**, Visual azimuth directions of looming stimuli from −72°, −36°, 0°, 36°, and 72° (left to right). Traces below indicate the time course of stimulus size. **b**, Normalized GCaMP7f fluorescence for DNae014 (left panels, green, n = 9), and DNb01 (right panels, magenta, n = 12) in response to each stimulus direction for left cell (L), right cell (R), and bilateral differential (black, R-L). Solid lines represent the grand mean of the mean traces for individual flies, the shaded areas indicated the boot-strapped 95% confidence intervals (CIs). **c**, Wingstroke amplitude responses to every stimulus direction of left (L), right (R), and differential (L-R) wingstroke amplitudes. **d**, Bilateral difference (R-L) ΔF/F for peak stimulus responses to each loom direction, plotted as mean of means with boot-strapped 95% CIs. DNae014 (green) responded more strongly but with a similar sign relative to stimulus direction, compared to DNb01 (magenta). Peak responses for DNae014 ΔF/F were larger than those for DNb01 (p < 0.001; two-way ANOVA with cell type and loom direction as the main effects and individual as a random effect).

**Extended Data Fig. 3.**
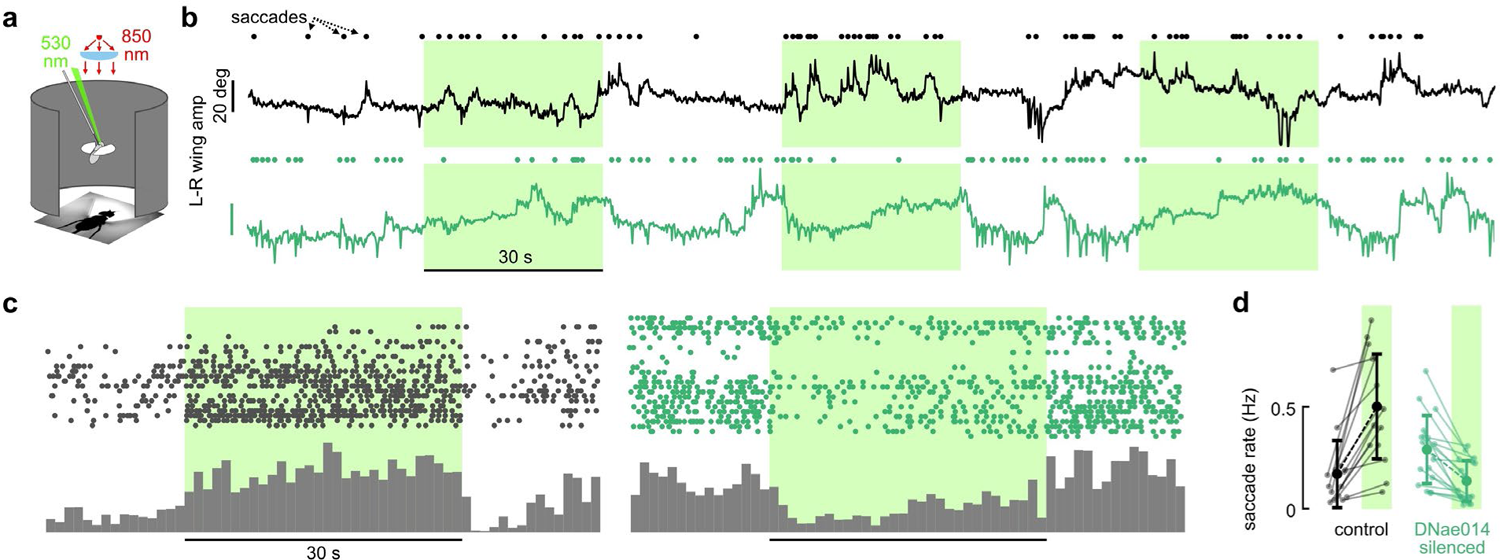
Temporal silencing of DNae014 reduces saccade rate. **a**, Tethered, flying flies expressing GtACR1 under split-GAL4 with exposure to 530 nm light (green shading). Wingstroke amplitudes (WSAs) were measured using a machine vision system with infra-red backlighting. The green light was focused on a 0.5 mm diameter spot on the posterior side of the head. **b**, Transient changes in L-R WSAs indicate fictive saccades. Saccades were automatically detected in 90 s flight trials by thresholding L-R WSA rate changes. An example empty vector control fly increased its spontaneous saccade rates during 30 s of 530 nm light exposure. An example fly expressing GtACR1 in DNae014 exhibited a reduction in saccade rate under green light exposure. **c**, Histograms of saccades pooled in 1-second-wide bins from 22 control flies showed an increased saccade rate during light exposures. In contrast, 25 flies expressing GtACR1 in DNae014 exhibited a reduced saccade rate during light exposure. **d**, Empty vector control flies saccaded at higher rates during green light exposure, whereas flies expressing GtACR1 in DNae014 saccaded at lower rates.

**Extended Data Fig. 4.**
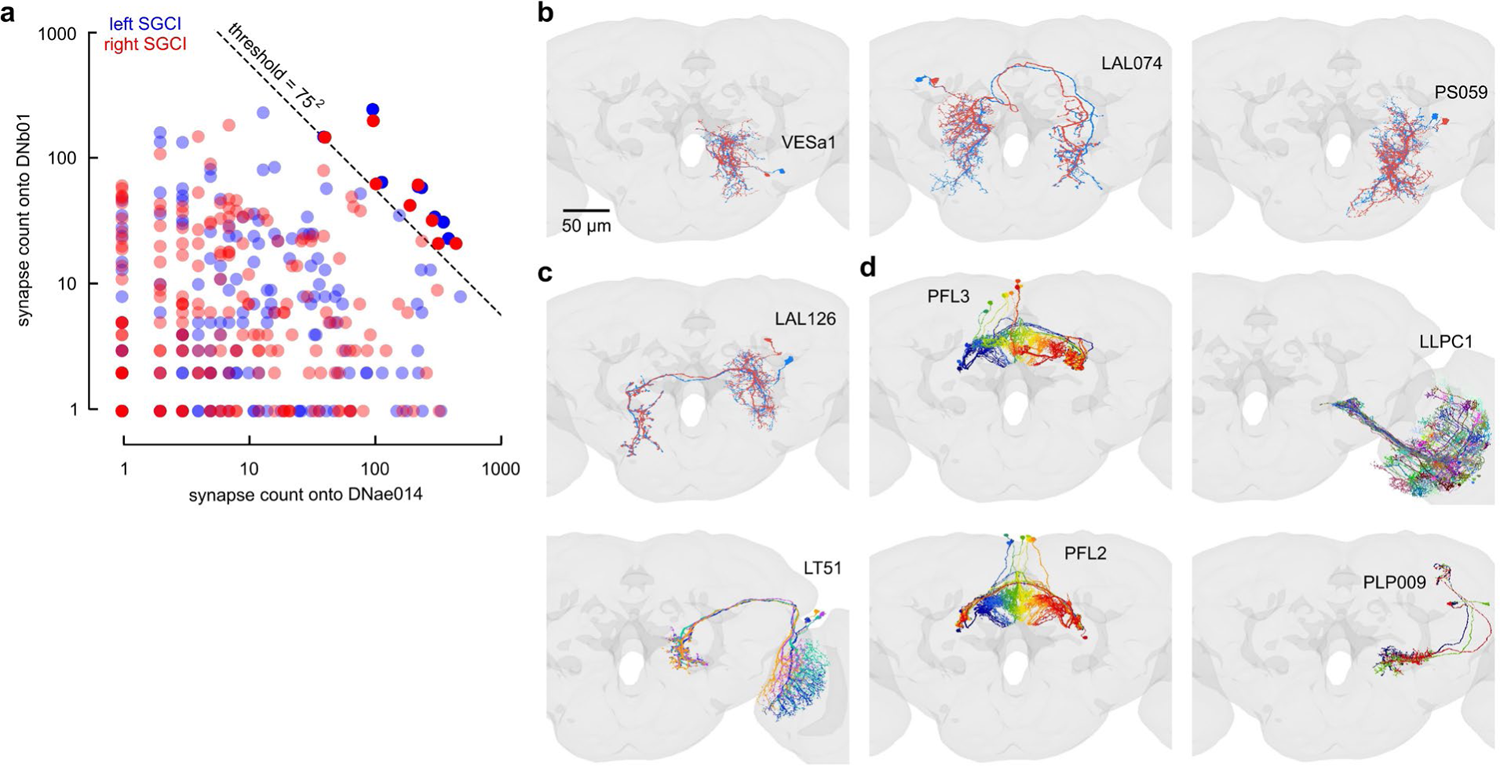
Identification of neuron types associated with saccade-generating couplets in FAFB. To identify shared inputs to saccade generating couplets (SGCs), neurons that form input connections to both unilateral DNae014 and DNb01 cells in a SGC were consolidated by type and subsequently thresholded based on combined synapse count onto both of the DN cells in a SGC. **a,** Candidate saccade-generating couplet input neurons (SGCIs, opaque circles in upper right corner of plot) were selected from the FAFB-Flywire dataset based on the criterion that the product of their synapse counts onto ipsilateral DNae014 and DNb01 neurons exceeded 75^2^ (= 5625) (dashed black line); this was the lowest threshold that preserved mirror-symmetry with respect to selection of left (blue) and right (red) SGCIs. **b-d,** Neurons were assigned to types based on similarities in morphology and connectivity in the FAFB and/or Hemibrain datasets. Colors indicate individual cells within a type. Symmetrical counterparts associated with the left SGC are not depicted for simplicity. **b,** Right SGCI neurons (VESa1, LAL074, PS059) that comprise types with more than bilateral pair of neurons. **c**, LAL126 neurons, which connect right SGIs to left SGCIs, similar to **b**. **d**, Neuron types (PFL3, LLPC1, LT51, PFL2, PLP009) that collectivity rank in the top ten types that synapse most heavily to one or more of the right SGCIs.

**Extended data Fig. 5.**
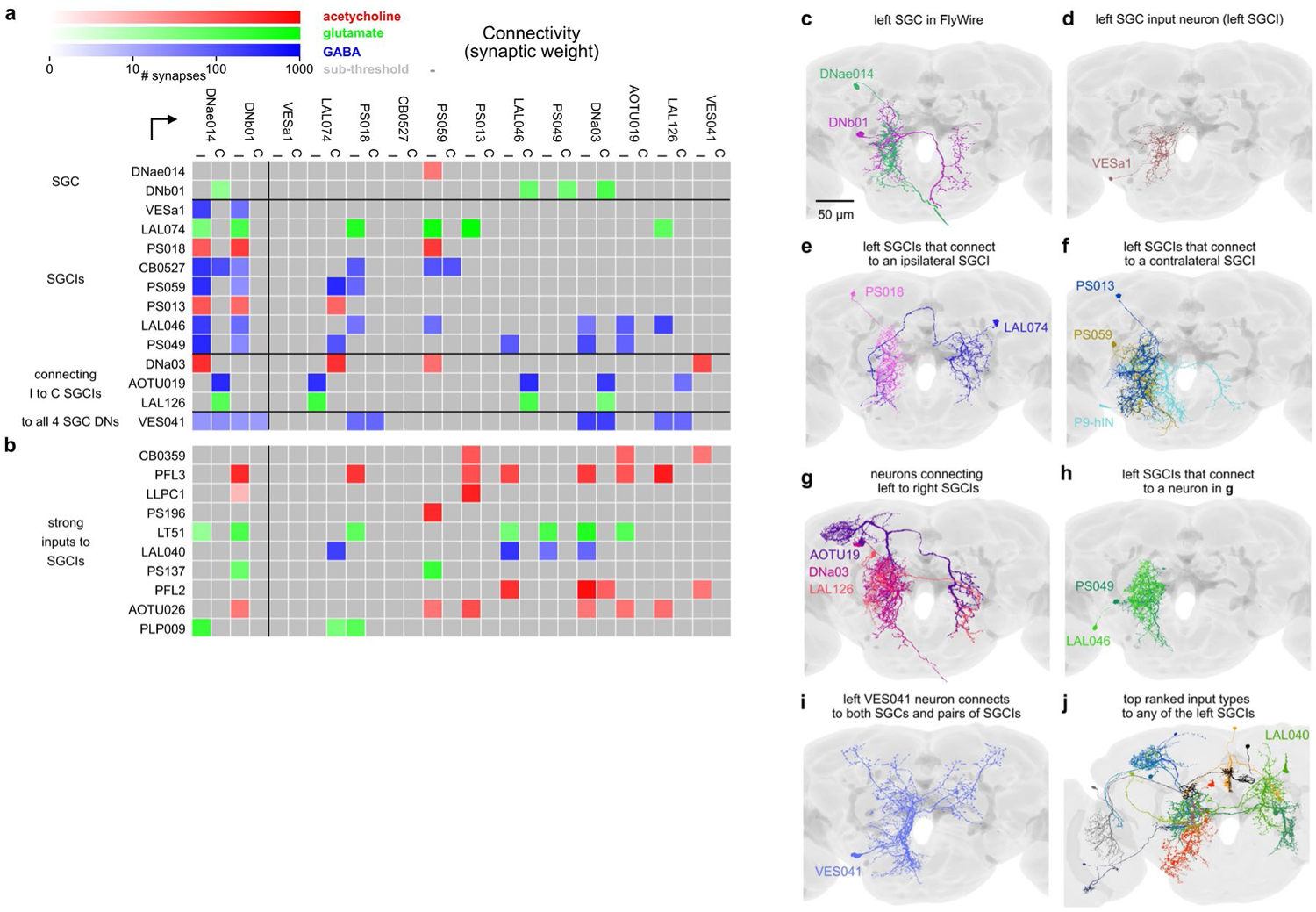
Unique connectivity of VES041 to DNae014 and DNb01 and their associated network partners. The table summarizes a quantitative analysis of the FAFB-Flywire dataset, in which we identified putative network partners of the SGCs. **a**, In addition to the DNae014 and DNb01 cells themselves, rows in the matrix represent neurons selected based on network topology and connectivity thresholding to form input synapses to both SGC cells (SGCIs), neuron types that form connections between ipsilateral and contralateral SGCIs (I to C SGCIs), and a neuron that connects to all four SGC cells (VES041). Columns indicate ipsilateral (I) and contralateral (C) pairs of these neurons. Matrix values are color-coded based on predicted transmitter classification and synaptic strength. VES041 is the only cell making synaptic connections with both ipsilateral and contralateral DNae014 and DNb01 cells and several upstream network partners. **b**, Top ten neuron types with the highest synapse count onto SGCIs, similar to **a**. Machine learning algorithms trained on the FAFB database suggest VES041 provides inhibitory, GABAergic input (blue), whereas DNae014 is cholinergic (red), and DNb01 glutamatergic (green). **c**-**j**, Morphologies in FAFB of neuron type examples cells listed in **a** and **b**. **c**, Left SGC consisting of DNae014 (green) and DNb01 (magenta) cells (from FAFB database). **d**, Left VESa1, a neuron connected to the left SGC. **e**, PS018 and LAL074, two SGCIs with connections to an ipsilateral SGCI. **f**, PS013, PS059, and P9-hIN, three SGCIs with connections to a contralateral SGCI. **g**, AOTU19, DNa03, and LAL126, three neurons that connect left and right SGCIs. **h**, Left SGCIs connected to an intermediary neuron. **i**, Left VES041 cell connects to all four cells in the saccade couplets. **i**, **j**, Top ten neuron types with most synaptic connections to the putative saccade network.

**Extended data Fig. 6.**
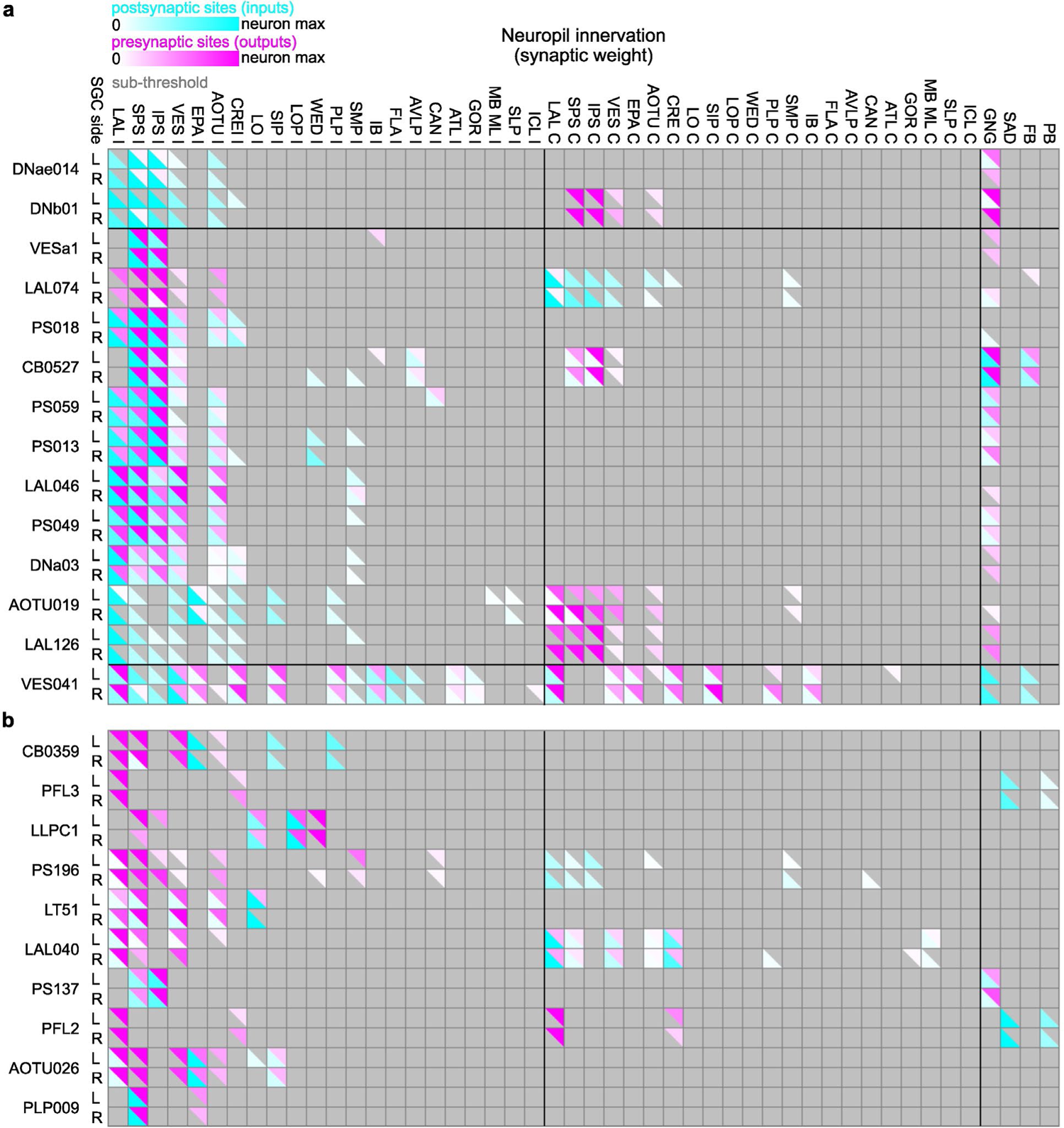
Summary of brain regions providing input to or receiving input from neurons implicated in generation and regulation of flight saccades. To coarsely assess where relevant neurons receive and send information, we present the predominant neuropils of the brain innervated by neurons related to the putative saccade network. a, Brain regions (columns) with postsynaptic dendrites (cyan) or presynaptic axon terminals (magenta) with over 75 synapses of SGC descending neurons and their putative input neuron types. Color tint represents relative synapse count, normalized per neuron. Type abbreviations are from the FAFB-Flywire dataset (Schlegel et al., 2023); brain regions from (Ito et al., 2014). b, Top ten neuron types with the highest synapse count onto putative network neurons, similar to a.

**Extended Data Fig. 7.**
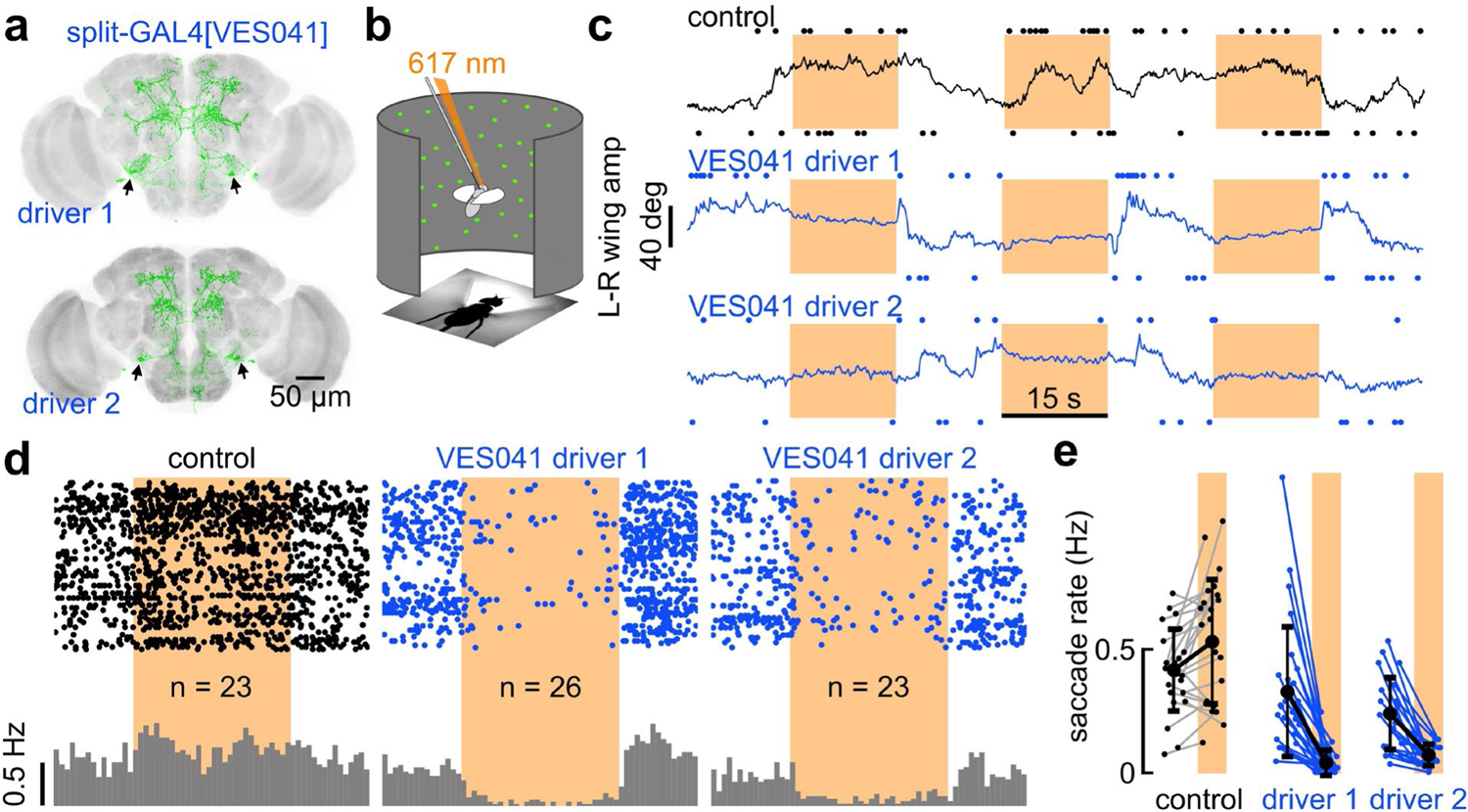
Activation of GABAergic VES041 neurons suppresses spontaneous saccades. **a**, Two split-GAL4 lines constructed to target the VES041 neuron (green, GFP). **b, c** Optogenetic activation of VES041 in a rigid tether arena. A focused LED light provided three 15-seconds periods of 623 nm excitation to the posterior part of the head capsule (orange beam in **b**, orange patches in **c**). **c,** Top: Empty vector control fly, with right and left saccades identified using the L-R WSA (dots above and below the panel, and black trace, respectively). Middle and bottom: Activation of VES041 drivers 1 and 2 (in blue). **d,** Each set of three rows represents orange light exposures for an individual fly (controls, left; VES041 split-GAL4 drivers 1 and 2 activations, middle and right). Gray histograms indicate mean saccade rate across populations. **e,** Individual saccade rates during light off and light on periods (orange patches) in control and VES041 drivers 1 and 2 groups; Each line represents an individual fly (n = 23, 26, and 23).

**Extended Data Fig. 8.**
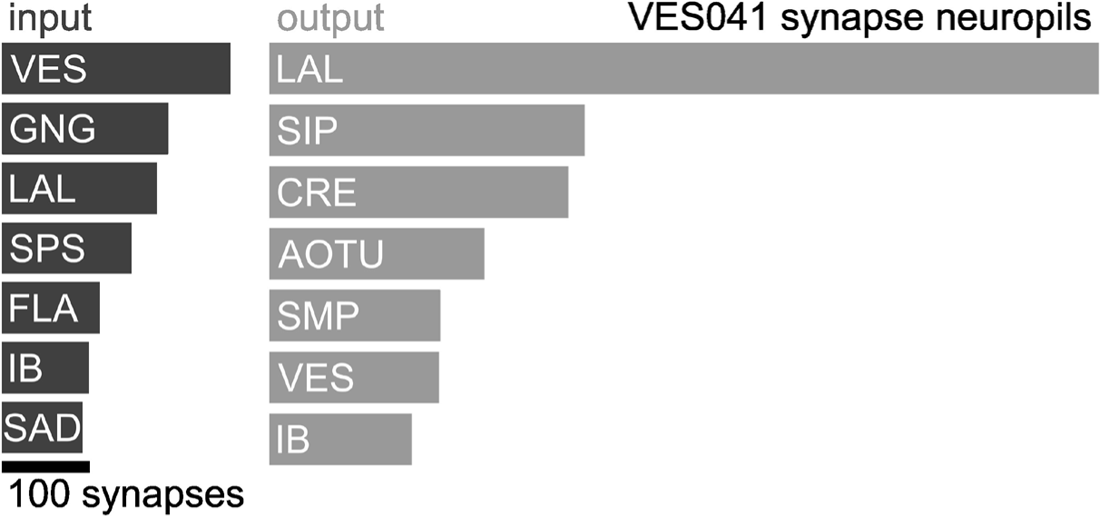
VES041 receives input from brain regions associated with sensory modalities and multi-modal integration. Top neuropils with highest averaged input (left column) and output synapse counts (right column) for the two VES041 cells in the FAFB dataset. Abbreviations: VES, vest; GNG, gnathal ganglia; LAL, lateral accessory lobes; SPS, superior posterior slope; FLA, flange; IB, inferior bridge; SAD, saddle; SIP, superior intermediate protocerebrum; CRE, crepine; AOTU, anterior optic tubercle; SMP, superior medial protocerebrum.

## Extended Data tables

**Extended Data Table 1.** Neurotransmitter predictions for DNae014, DNb01, and VES041.

|  | left<br>DNae014 | right<br>DNae014 | left<br>DNb01 | right<br>DNb01 | left<br>VES041 | right<br>VES041 |
| --- | --- | --- | --- | --- | --- | --- |
| GABA | 0.05 | 0.06 | 0.08 | 0.06 | **0.86** | **0.84** |
| acetylcholine | **0.60** | **0.71** | 0.13 | 0.06 | 0.08 | 0.08 |
| glutamate | 0.25 | 0.13 | **0.76** | **0.87** | 0.05 | 0.07 |
| octopamine | 0.01 | 0.00 | 0.00 | 0.00 | 0.00 | 0.00 |
| serotonin | 0.06 | 0.05 | 0.03 | 0.00 | 0.00 | 0.00 |
| dopamine | 0.04 | 0.05 | 0.01 | 0.00 | 0.00 | 0.00 |

Values are confidence scores for neurotransmitter predictions.

**Extended Data Table 2.** Interconnectivity between DNs.

|  | left<br>DNae014 | right<br>DNae014 | left<br>DNb01 | right<br>DNb01 |
| --- | --- | --- | --- | --- |
| left DNae014 | - | 0 | 2 | 0 |
| right DNae014 | 0 | - | 0 | 7 |
| left DNb01 | 0 | **21** | - | 0 |
| right DNb01 | **18** | 0 | 0 | - |

Values are synaptic weight.

**Extended Data Table 3.** List of neurons used in the connectivity analysis.

| root ID | MS type | codex name | schlegel type | hemibrain type |
| --- | --- | --- | --- | --- |
| 720575940609488942 | DNae014 | DNa.24 | DNae014 | PS017 |
| 720575940631759663 | DNae014 | DNa.18 | DNae014 | PS017 |
| 720575940630758418 | DNb01 | DNb.6 | DNb01 | DNb01 |
| 720575940625878160 | DNb01 | DNb.4 | DNb01 | DNb01 |
| 720575940611859570 | VESa1 | SPS.IPS.14 | NA | NA |
| 720575940613949090 | VESa1 | SPS.IPS.12 | NA | NA |
| 720575940610727630 | VESa1 | SPS.IPS.19 | NA | NA |
| 720575940619358800 | VESa1 | SPS.IPS.15 | NA | NA |
| 720575940616807897 | LAL074 | LAL074.4 | LAL074, LAL084 | LAL074, LAL084 |
| 720575940638681917 | LAL074 | LAL074.3 | LAL074, LAL084 | LAL074, LAL084 |
| 720575940619866027 | LAL074 | LAL074.1 | LAL074, LAL084 | LAL074, LAL084 |
| 720575940617226141 | LAL074 | LAL074.2 | LAL074, LAL084 | LAL074, LAL084 |
| 720575940617100985 | PS018 | PS018b.2 | PS018b | PS018 |
| 720575940605495072 | PS018 | LAL.IPS.10 | PS018b | PS018 |
| 720575940617444130 | CB0527 | suppressorofDNs1.1 | CB0527 | NA |
| 720575940623455239 | CB0527 | suppressorofDNs1.2 | CB0527 | NA |
| 720575940619681006 | PS059 | PS059.1 | PS059 | PS059 |
| 720575940631721785 | PS059 | PS059.2 | PS059 | PS059 |
| 720575940614570415 | PS059 | PS059.4 | PS059 | PS059 |
| 720575940619549598 | PS059 | PS059.3 | PS059 | PS059 |
| 720575940607982428 | PS013 | PS013.1 | PS013 | PS013 |
| 720575940620842111 | PS013 | PS013.2 | PS013 | PS013 |
| 720575940635970190 | LAL046 | LAL046.1 | LAL046 | LAL046 |
| 720575940617243393 | LAL046 | LAL046.2 | LAL046 | LAL046 |
| 720575940618058048 | PS049 | PS049.1 | PS049 | PS049 |
| 720575940623533749 | PS049 | PS049.2 | PS049 | PS049 |
| 720575940630085583 | DNa03 | DNa03.1 | DNa03 | DNa03 |
| 720575940620918789 | DNa03 | LAL.IPS.1 | DNa03 | DNa03 |
| 720575940631517251 | AOTU019 | AOTU019.2 | AOTU019 | AOTU019 |
| 720575940627100584 | AOTU019 | AOTU019.1 | AOTU019 | AOTU019 |
| 720575940630040438 | LAL126 | LAL126.2 | LAL126 | LAL126 |
| 720575940625054906 | LAL126 | LAL.IPS.22 | LAL126 | LAL126 |
| 720575940628290307 | LAL126 | LAL126.1 | LAL126 | LAL126 |
| 720575940626717573 | LAL126 | LAL.IPS.21 | LAL126 | LAL126 |
| 720575940635202204 | VES041 | VES041.2 | VES041 | VES041 |
| 720575940627253604 | VES041 | VES041.1 | VES041 | VES041 |
| 720575940629883971 | CB0359 | AOTU016.2 | CB0007 | NA |
| 720575940623650055 | CB0359 | AOTU016.3 | CB0359 | NA |
| 720575940623430621 | PFL3 | FB.LAL.15 | PFL3 | PFL3 |
| 720575940619006087 | PFL3 | PFL3.4 | PFL3 | PFL3 |
| 720575940628692841 | LLPC1 | Lccn2.17 | LLPC1 | LLPC1 |
| 720575940623097271 | LLPC1 | LLPC2a.12 | LLPC1 | LLPC1 |
| 720575940625082969 | PS196 | PS196.1 | PS196 | PS196 |
| 720575940629229020 | PS196 | PS196.2 | PS196 | PS196 |
| 720575940617260374 | LT51 | LT51.8 | LT51 | LT51 |
| 720575940639563728 | LT51 | LT51.12 | LT51 | LT51 |
| 720575940622018217 | LAL040 | LAL040.1 | LAL040 | LAL040 |
| 720575940617818708 | LAL040 | LAL040.2 | LAL040 | LAL040 |
| 720575940624447240 | PS137 | PS137.2 | PS137 | PS137 |
| 720575940633333017 | PS137 | PS137.4 | PS137 | PS137 |
| 720575940635467150 | PFL2 | PFL2(PB12b).4 | PFL2 | PFL2 |
| 720575940623937976 | PFL2 | PFL2(PB12b).5 | PFL2 | PFL2 |
| 720575940628916444 | AOTU026 | AOTU.LAL.8 | AOTU026 | AOTU026 |
| 720575940619738863 | AOTU026 | AOTU.LAL.6 | AOTU026 | AOTU026 |
| 720575940624388878 | PLP009 | SPS.160 | PLP009 | PLP009 |
| 720575940618540081 | PLP009 | SPS.138 | PLP009 | PLP009 |

## Extended Data video captions

**Extended Data Vid. 1 |** Expression pattern of the split-GAL4 driver line for DNae014, w[1118]; P{y[+t7.7] w[+mC]=VT025718-p65.AD}attP40/CyO; P{y[+t7.7] w[+mC]=GMR56G08.DBD}attP2. As in all following videos, the optical sections appear from posterior to anterior in the brain, and from dorsal to ventral in the ventral nerve cord. GFP expression is shown in green; nc82 staining in magenta. To reduce file size, images were compressed using MPEG-4. This driver line was generated in our lab.

**Extended Data Vid. 2 |** Expression pattern in the ventral nerve cord of the split-GAL4 driver line for DNae014, similar to Extended Data Video 1.

**Extended Data Vid. 3 |** Expression pattern in the brain of the split-GAL4 driver line for DNb01, SS02383 (P{w[+mC]=BJD103G04-pBPp65ADZpUw}attP40; P{w[+mC]=GMR45H03-pBPZpGdbdUw}attP2). The driver line was generated at Janelia Research Campus.

**Extended Data Vid. 4 |** Expression pattern in the ventral nerve cord of the split-GAL4 driver line for DNb01, similar to Extended Data Video 3.

**Extended Data Vid. 5 |** Expression pattern in the brain of the split-GAL4 driver line 1 for VES041, w[1118]; P{y[+t7.7] w[+mC]=GMR64G09-p65.AD}attP40/CyO; P{y[+t7.7] w[+mC]=VT044519.DBD}attP2/TM6B. CsChrimson-mVenus expression is shown in green; nc82 staining in magenta. This driver line was generated in our lab.

**Extended Data Vid. 6 |** Expression pattern in the ventral nerve cord of the split-GAL4 driver line 1 for VES041, similar to Extended Data Video 5.

**Extended Data Vid. 7 |** Expression pattern in the brain of the split-GAL4 driver line 2 for VES041, w[1118]; P{y[+t7.7] w[+mC]=GMR64G09-p65.AD}attP40/CyO; P{y[+t7.7] w[+mC]=GMR31E10.DBD}attP2/TM6B. CsChrimson-mVenus expression is shown in green; nc82 staining in magenta. This driver line was generated in our lab.

**Extended Data Vid. 8 |** Expression pattern in the ventral nerve cord of the split-GAL4 driver line 2 for VES041, similar to Extended Data Video 7.

## Notes

### Competing Interest Statement

The authors have declared no competing interest.

### Summary of Updates

The manuscript has been revised at the suggestions of colleagues to include additional references regarding the creation of split-GAL4 lines and connectomics methods.

